# R-loops orchestrate the maternal-to-zygotic transition by harnessing RNA polymerase II pause release

**DOI:** 10.1101/2025.11.03.686253

**Authors:** Yaoyi Li, Qing Li, Xinxiu Wang, Chao Di, Yingliang Sheng, Qingqing Cai, Sainan Huang, Jiayu Chen, Guangming Wu, Shaorong Gao, Hongjie Yao

## Abstract

R-loops, ubiquitous three-stranded nucleic acid structures abundant in the mammalian genome, play pivotal roles in diverse biological processes^1–3^. However, the dynamics and functions of R-loops during mammalian preimplantation embryonic development remain poorly understood. Here we optimized an innovative, ultra-sensitive RIAN-seq method capable of mapping genome-wide R-loop landscape using ultra-low cell numbers to investigate R-loop dynamics in mouse gametes and early embryos. Our findings reveal the widespread presence of AT-rich R-loops during early mouse embryonic development and highlight their critical roles in orchestrating transcription machinery during zygotic genome activation (ZGA). We demonstrate that the stability and inheritance of R-loops through developmental stages are governed by their sequence composition, with GC-rich R-loops exhibiting greater stability. Notably, R-loops, particularly AT-rich R-loops, inhibit DDX21 helicase activity on the 7SK/HEXIM1 snRNP complex, thereby reducing CDK9 release. This, in turn, modulates the phosphorylation of Ser2 at C-terminal domain (CTD) of RNA polymerase II (RNAPIIS2p) at the promoters of major ZGA and maternal genes. Furthermore, R-loops promote RNAPII accumulation at major ZGA gene promoters, preventing RNAPII from premature release and ensuring timely activation of major ZGA genes. Conversely, R-loops enhance the retention of stalled RNAPII at maternal gene promoters, effectively repressing their expression and facilitating the maternal-to-zygotic transition. In summary, our study unveils a key role for R-loops in regulating the maternal-to-zygotic transcription transition and early embryonic development in mice.

## Main

R-loops, composed of an RNA:DNA hybrid and a displaced single-stranded DNA, are widespread across the genome and closely associated with transcription^4–6^, as well as DNA replication and repair processes^7^. While scheduled R-loops play essential roles in transcription termination^1,8^ and gene expression^9,10^, aberrant R-loop accumulation or dysregulation of R-loop regulatory factors has been implicated in disease^11^, abnormal development^12^ and disruption of cellular homeostasis^13^. Despite their well-established roles in these fundamental processes, the functional significance of R-loops during mouse preimplantation embryonic development and their potential involvement in transcriptional regulation remain poorly understood, representing a significant gap in our knowledge of R-loop biology.

Here, we leveraged our recently developed RIAN-seq (R-loop identification assisted by nucleases and sequencing)^14^, optimized its protocol to achieve a highly sensitive and efficient method for low-cell input material, enabling comprehensive mapping of R-loop dynamics during mouse preimplantation embryonic development. Our findings reveal that R-loops undergo extensive reprogramming and play a crucial role in regulating RNA polymerase II (RNAPII) transcription during early embryogenesis. Notably, the loss of R-loops, particularly AT-rich R-loops, leads to significant downregulation of major ZGA genes and the aberrant persistence of maternal transcripts, ultimately resulting in disrupted embryonic development.

### Genome-wide landscape of R-loop dynamics in mouse gametes and preimplantation embryos

To investigate the dynamic landscape of R-loops during mouse preimplantation embryonic development, we implemented our optimized RIAN-seq method and established a streamlined one-tube workflow (Fig. 1a), incorporating AtacWorks^15^-based denoising, specifically tailored for limited input materials (Extended Data Fig. 1a-c). Method validation using HEK293T cells demonstrated that RIAN-seq profiles from low-input-cell samples (10^3^, 10^2^, 10 cells) faithfully recapitulated those from bulk-cell samples (Extended Data Fig. 1d-f). Furthermore, base composition analysis confirmed comparable GC/AT content of R-loops between low-input and bulk-cell samples (Extended Data Fig. 1g), establishing RIAN-seq as a robust platform for R-loop profiling in scarce biological samples.

**Fig. 1.**
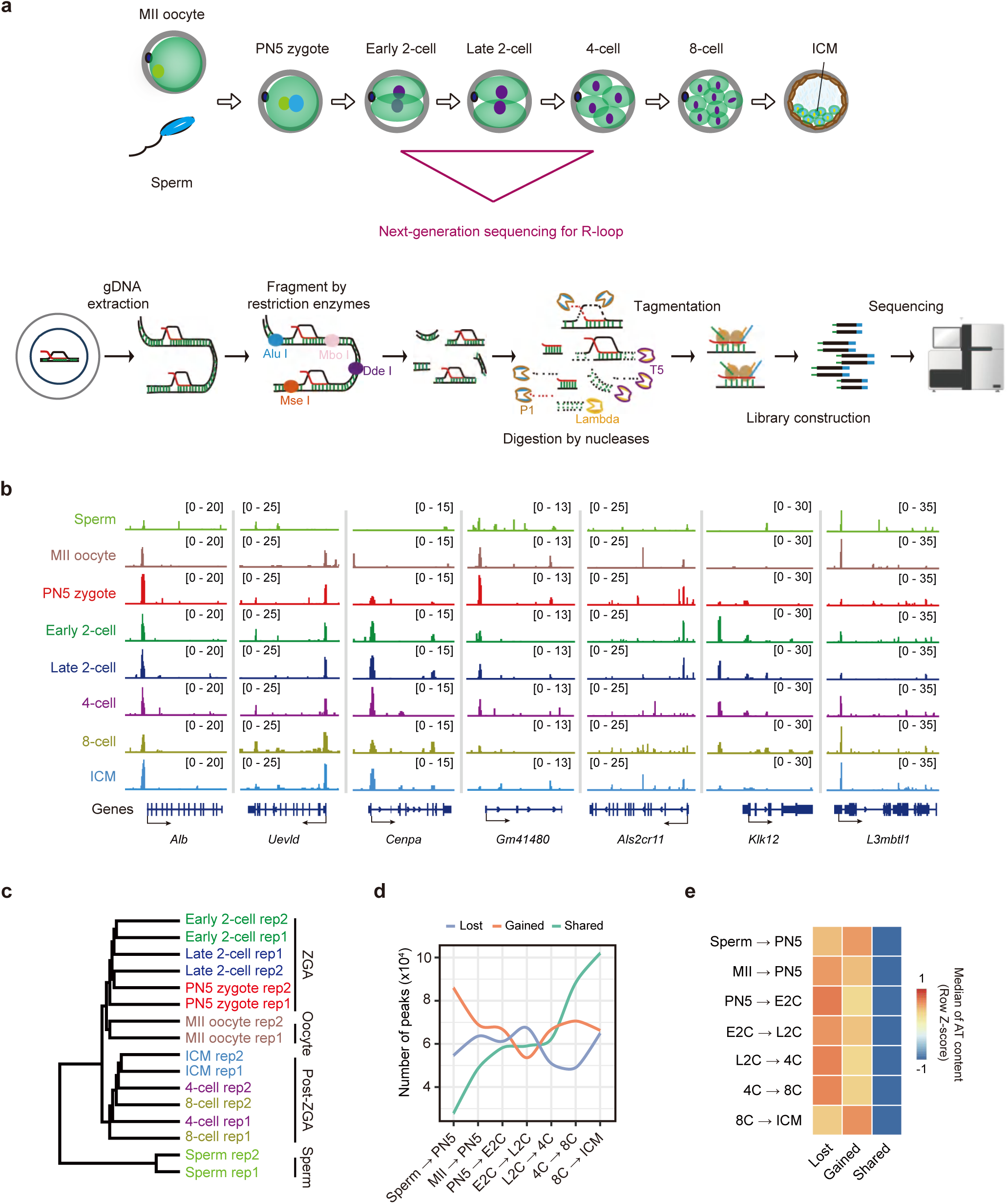
Genome-wide profiles of R-loops in mouse gametes and early preimplantation embryos. **a**, Schematic representation of mouse gametes and early preimplantation embryos subjected for R-loop analysis by RIAN-seq. The restriction enzymes used include: Alu I, DdeI, Mbo I, Mse I. P1, nuclease P1; Lambda, Lambda exonuclease; T5, T5 exonuclease. **b**, Tracks showing R-loop signals in mouse gametes and early preimplantation embryos. **c**, Hierarchical clustering analysis illustrating global R-loop enrichment across biological replicates in mouse gametes and early preimplantation embryos (two biological replicates, Rep). **d**, Curve graphs showing the quantification of gained, lost, and shared R-loops between consecutive developmental stages of mouse gametes and early preimplantation embryos. MII, MII oocyte; PN5, PN5 zygote; E2C, early 2-cell; L2C, late 2-cell; 4C, 4-cell; 8C, 8-cell; ICM, inner cell mass from the blastocyst. **e**, Heatmaps depicting the AT content of gained, lost, and shared R-loops between adjacent stages of mouse gametes and early preimplantation embryos. Data are presented as median values.

Immunostaining revealed authentic R-loop signals occurred across MII oocytes and early embryos (Extended Data Fig. 2a,b). Using RIAN-seq, we next investigated R-loop dynamics in mouse gametes and preimplantation embryos. RIAN-seq identified R-loops in sperm, MII oocytes and early preimplantation embryos, spanning from PN5 zygotes to inner cell masses (ICM) of blastocysts (Fig. 1a,b), with high reproducibility between two biological replicates (R > 0.9) (Extended Data Fig. 2c,d). The specificity of R-loops captured by RIAN-seq was validated by a dramatic reduction in signals following RNase H treatment (Extended Data Fig. 2e-g). Immunostaining also observed R-loop signals at transcriptionally quiescent MII oocytes, which were effectively resolved by RNase H treatment (Extended Data Fig. 2a,b), suggesting that R-loops form *in trans*. Notably, from 4-cell stage onward, R-loop signals at promoters showed a stronger correlation with transcriptional output levels (Extended Data Fig. 2h), aligning with the well-established positive association between R-loops and transcription^2,3^. Collectively, these findings demonstrate that RIAN-seq generates high-quality and reliable R-loop signals from mouse gametes and early preimplantation embryos.

R-loops were highly enriched at promoters in mouse gametes and early preimplantation embryos (Fig. 1b and Extended Data Fig. 3a), with both GC-and AT-skewed sequences present within R-loop regions (Extended Data Fig. 3b). Quantitative analysis revealed a reduction in both the numbers and genomic coverage of R-loops during ZGA stages compared to other stages of preimplantation development (Extended Data Fig. 3c,d). The majority of R-loops in mouse gametes and embryos were approximately 100 bp in length (Extended Data Fig. 3e), with no substantial differences in their genomic distribution between gametes and preimplantation embryos (Extended Data Fig. 3f). Approximately 50% of R-loops were localized to protein-coding genes, while 25% were associated with transposon repeats (Extended Data Fig. 3g). Hierarchical clustering analysis identified fertilization and ZGA as key time points for the remodeling of R-loop dynamics (Fig. 1c). Stage transition analysis revealed that the most extensive R-loop remodeling occurred during the gamete-to-zygote transition, whereas changes during ZGA were more subtle (Fig. 1d). Shared R-loops progressively accumulated, particularly after late 2-cell stage (Fig. 1d and Extended Data Fig. 4a). Sequence analysis showed that shared R-loops preferentially formed at GC-rich regions, whereas gained and lost R-loops were predominantly AT-rich (Fig. 1e), suggesting that AT-rich R-loops exhibit stage-specific formation. Quantitative analysis further revealed a progressive decline in stage-specific R-loops during ZGA (Extended Data Fig. 4b), with 73.06% to 76.7% of stage-specific R-loops preferentially enriched at AT-rich regions in mouse gametes and early embryos (Extended Data Fig. 4c,d). These findings highlight the highly dynamic nature of AT-rich R-loops during early embryonic development.

To explore the potential roles of stage-specific R-loops, we performed Gene Ontology (GO) analysis and found that R-loops specific to early developmental stages (from PN5 zygote to late 2-cell stage) were enriched in genes associated with translation and histone modifications (Extended Data Fig. 4e). R-loops predominantly enriched at the late stages (from 4-cell to ICM stage) were closely associated with genes involved in cell fate commitment and embryo implantation (Extended Data Fig. 4e). These results suggest that the dynamic reprogramming of R-loops may play a crucial role in regulating key processes during mouse preimplantation embryonic development.

In previous report on cell lines, unmethylated CpG island promoters were found to be enriched with R-loops^16^. We then categorized all promoters into high, intermediate, and low CpG promoters (HCPs, ICPs, and LCPs) and observed a positive correlation between R-loop levels and promoter GC content in mouse gametes and preimplantation embryos (Extended Data Fig. 4f). Notably, genomic regions with higher R-loop levels exhibited lower DNA methylation in both MII oocytes and preimplantation embryos (Extended Data Fig. 4g), and genome-wide analysis demonstrated a negative correlation between R-loop signals and DNA methylation levels in mouse gametes and preimplantation embryos (Extended Data Fig. 4h). Furthermore, R-loop-marked regions in mouse gametes underwent significant demethylation following fertilization and maintained low methylation levels from the 2-cell stage onward (Extended Data Fig. 4i). These findings align with previous report that R-loops recruit demethylation machinery to sustain DNA hypomethylation^17^.

Collectively, these findings demonstrate extensive R-loop remodeling following fertilization and major ZGA, with R-loop dynamics being profoundly shaped by underlying sequence composition and developmental context.

### Stage-specific enrichment of AT-rich R-loops during zygotic genome activation

Given the extensive reprogramming of genome-wide R-loops following fertilization and major ZGA, we examined the dynamics of R-loops specifically during ZGA stages. Prior to major ZGA, R-loops exhibited minimal association with actively transcribing genes; however, their association with transcribing genes progressively increased from late 2-cell to blastocyst stages (Fig. 2a). Notably, R-loop levels at the promoters of both constantly active and inactive genes were lower at the ZGA stages after fertilization (Fig. 2b and Extended Data Fig. 5a), suggesting a potential RNAPII-independent mechanism of R-loop formation during this critical developmental stage. Furthermore, we observed the low expression levels of RNase H family members, key R-loop resolving factors^10,18^, during ZGA (Extended Data Fig. 5b), suggesting that the reduced R-loop levels were not attributable to increased RNase H-mediated resolution.

**Fig. 2.**
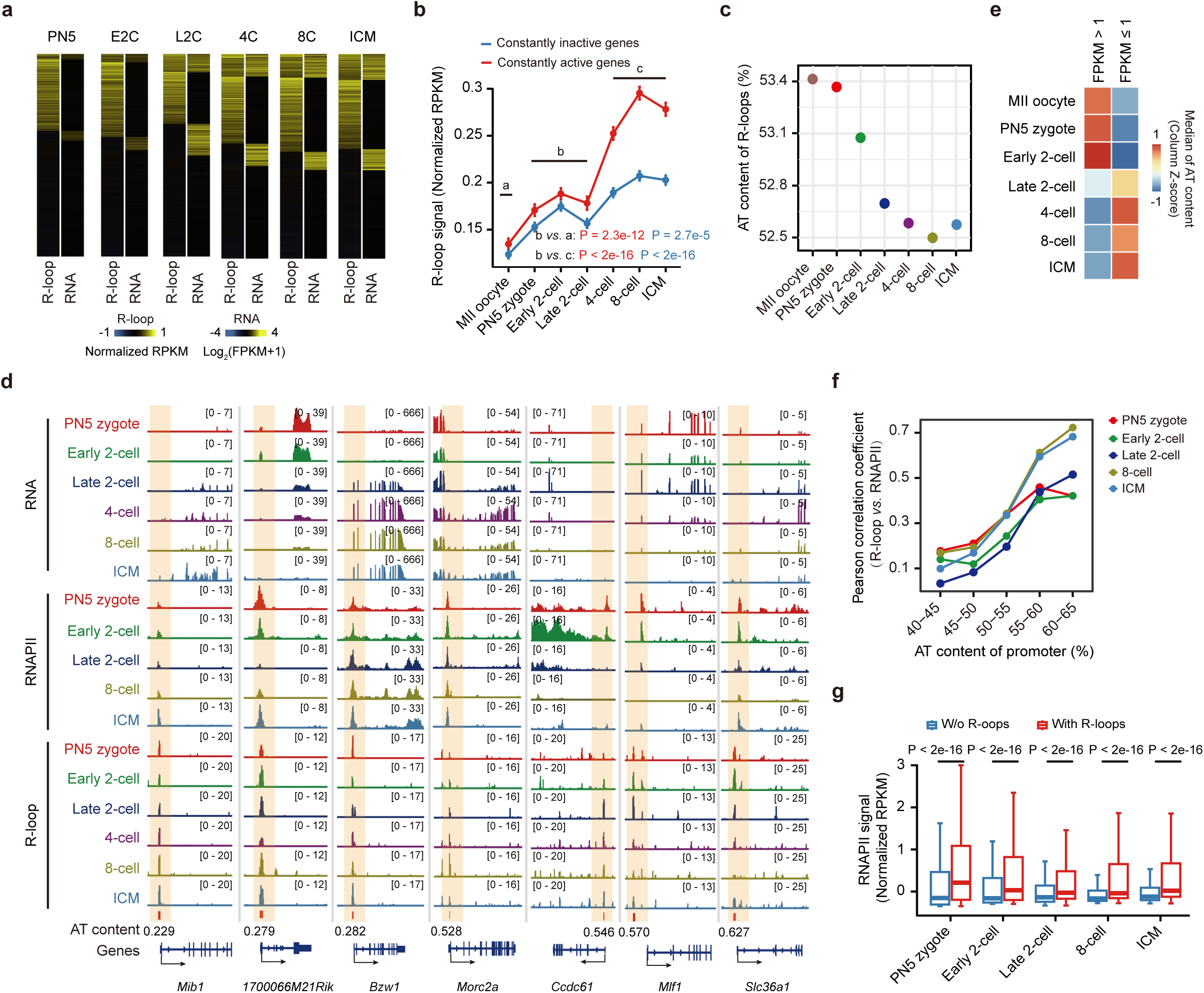
AT-rich R-loops are abundant during early embryonic development. **a**, Heatmaps showing the expression of genes (MII FPKM < 0.5) and R-loop enrichment (Z-score normalized) at their promoters. FPKM, fragments per kilobase of transcript per million mapped reads; RPKM, reads per kilobase of transcript per million mapped reads. PN5, PN5 zygote; E2C, early 2-cell; L2C, late 2-cell; 4C, 4-cell; 8C, 8-cell; ICM, inner cell mass from the blastocyst. **b**, Line charts showing promoter R-loop enrichment (Z-score normalized) of constantly active genes (red) and constantly inactive genes (blue) during early preimplantation embryos. Error bars, mean ± SE. a: Average R-loop signals in MII oocytes; b: Average R-loop signals at ZGA stages; c: Average R-loop signals at post-ZGA stages. P values, two-sided Wilcoxon rank-sum test. **c**, AT content analysis of R-loops in mouse MII oocytes and early preimplantation embryos. Error bars, mean ± SE. **d**, Tracks showing gene expression, R-loop, and RNAPII at representative loci with high or low AT content in early preimplantation embryos. Corresponding AT content is indicated, R-loop and RNAPII enrichments near TSSs are shaded. **e**, Heatmaps depicting AT content of the promoters for lowly expressed (FPKM ≤ 1) and highly expressed (FPKM > 1) genes in mouse MII oocytes and early preimplantation embryos. Data are presented as median values. **f**, Line charts illustrating the Pearson correlation between R-loop signals and RNAPII occupancy at the promoters categorized by AT content during early preimplantation embryos. **g**, Box plots showing RNAPII enrichment (Z-score normalized) at AT-rich promoters (AT content > 0.5) with or without R-loops in early preimplantation embryos. P values, two-sided Wilcoxon rank-sum test. Center line, median; box, 25th and 75th percentiles; whiskers, 1.5 × interquartile range (IQR).

Sequence analysis demonstrated higher AT content in R-loops during ZGA compared to other developmental stages (Fig. 2c, Extended Data Fig. 5c), with certain loci displaying particularly strong AT-rich R-loop signals (Fig. 2d). AT-rich R-loops exhibit greater dynamics and stage specificity during early embryonic development (Fig. 1d,e and Extended Data Fig. 4c,d), diverging from the predominant GC-rich R-loops in mammalian cell lines^19^, but mirroring the prevalence of AT-rich R-loops in the genomes of non-mammalian organisms, such as *Arabidopsis*^20^ and yeast^21^. Further investigation of promoter sequence features across different gene expression levels uncovered an intriguing pattern: GC-rich promoters were predominantly associated with lowly expressed genes during ZGA, whereas AT-rich promoters correlated with highly expressed genes (Fig. 2e). This pattern completely reversed after major ZGA (Fig. 2e), highlighting the stage-specific nature of R-loop regulation during development.

Despite the weak correlation between promoter R-loops and transcriptional output before late 2-cell stage (Fig. 2a and Extended Data Fig. 2h), the colocalization of promoter R-loops with RNAPII was observed during ZGA (Extended Data Fig. 5d). Notably, RNAPII signals positively correlated with R-loop levels at promoters, and this association strengthened with increasing AT content of promoters (Fig. 2f and Extended Data Fig. 5e). Additionally, genes with AT-rich promoters harboring R-loops exhibited higher RNAPII enrichment compared to those lacking R-loops (Fig. 2g), suggesting that a potential regulatory role for R-loops in orchestrating RNAPII configuration during early embryonic development.

### RNAPII-uncoupled R-loop formation at the promoters of major ZGA and maternal genes during maternal-to-zygotic transition

As RNAPII undergoes configuration in preparation for ZGA transcription^22^, we investigated the relationship between R-loops and RNAPII during ZGA. Specifically, we analyzed their interplay at the promoters of three distinct gene classes: maternal genes, major ZGA genes, and Polycomb group (PcG) target genes. Consistent with their AT-rich promoter composition (Extended Data Fig. 5f), major ZGA and maternal genes exhibited slightly weaker R-loop signals compared to PcG target genes, which possess GC-rich promoters (Fig. 3a and Extended Data Fig. 5f,g). Notably, the promoters of major ZGA and maternal genes displayed enhanced RNAPII enrichment (Extended Data Fig. 5h), with a higher degree of RNAPII pausing at these promoters during ZGA, compared to PcG target genes (Extended Data Fig. 5i,j). Additionally, we detected elevated R-loop and RNAPII levels at the promoters of major ZGA genes compared to other silent genes at PN5 zygote and early 2-cell stages (Fig. 3b,c). Both R-loop and RNAPII levels were higher at PN5 zygote and early 2-cell stages than at late 2-cell stage (Fig. 3c,d). Furthermore, R-loop levels showed a positive correlation with both promoter RNAPII enrichment and pausing index of major ZGA and maternal genes, with stronger association observed for major ZGA genes (Extended Data Fig. 5k,l). These data suggest a coordinated enrichment of R-loops and RNAPII at major ZGA genes during ZGA.

**Fig. 3.**
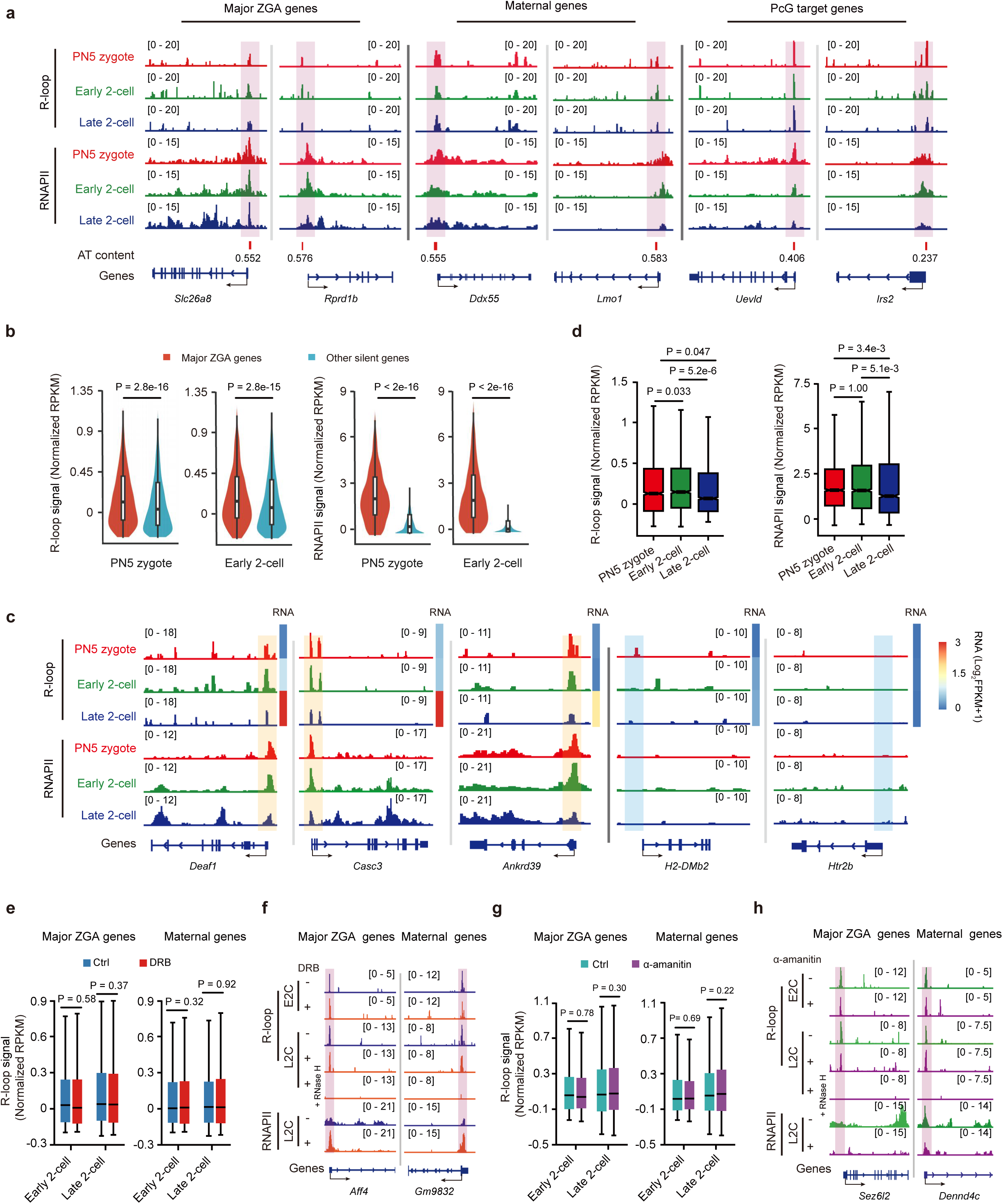
R-loops facilitate RNAPII accumulation prior to major ZGA. **a**, Tracks displaying promoter R-loop and RNAPII enrichment of major ZGA, maternal genes, and PcG target genes in embryos during ZGA. The AT content of R-loops is indicated. **b**, Violin plots comparing promoter R-loop and RNAPII enrichment (Z-score normalized) of major ZGA genes (n = 2,052) versus other transcriptionally silent genes (FPKM < 0.5, n = 10,985) at PN5 zygote and early 2-cell stages. **c**, Tracks and heatmaps illustrating R-loop, RNAPII enrichment and gene expression of representative genes during ZGA. R-loop and RNAPII near TSSs of major ZGA genes (orange) and other silent genes (blue) are shaded. **d**, Box plots quantifying promoter R-loop and RNAPII enrichment (Z-score normalized) of major ZGA genes (n = 2,052) during ZGA. **e**, Box plots showing promoter R-loop enrichment (Z-score normalized) of major ZGA and maternal genes in early and late 2-cell embryos with or without DRB treatment. **f**, Tracks displaying R-loop and RNAPII enrichment at representative loci in early (E2C) and late 2-cell (L2C) embryos treated with or without DRB. Additional tracks showing R-loop signals following RNase H digestion upon DRB treatment at late 2-cell stage. **g**, Box plot illustrating promoter R-loop enrichment (Z-score normalized) of major ZGA and maternal genes in early and late 2-cell embryos with or without α-amanitin treatment. **h**, Tracks depicting R-loop and RNAPII enrichment at representative loci in early and late 2-cell embryos treated with or without α-amanitin. Additional tracks displaying R-loop signals following RNase H digestion upon α-amanitin treatment at late 2-cell stage. P values, two-sided Wilcoxon rank-sum test. Center line, median; box, 25th and 75th percentiles; whiskers, 1.5 × IQR.

Since RNAPII pre-configuration requires minor ZGA^22^, to determine whether R-loop formation depends on RNAPII configuration during ZGA, we conducted genome-wide R-loop mapping in early 2-cell and late 2-cell embryos treated with or without RNAPII elongation inhibitor DRB (5,6-dichloro-1-β-D-ribofuranosyl-benzimidazole) from PN2 stage. While DRB treatment led to a moderate reduction in global R-loop signals (Extended Data Fig. 6a-c), it did not affect R-loop signals at the promoters of major ZGA and maternal genes (Fig. 3e,f), despite promoting RNAPII accumulation at these promoters (Extended Data Fig. 6d,e). These findings suggest that R-loops at the promoters of major ZGA and maternal genes during ZGA are not a direct byproduct of RNAPII transcription.

To further explore whether R-loops respond to RNAPII enrichment, we treated embryos with α-amanitin to induce RNAPII degradation. While α-amanitin treatment led to overall increase in global R-loop signals (Extended Data Fig. 6f-h), it had no impact on R-loop signals at the promoters of major ZGA and maternal genes in early 2-cell and late 2-cell embryos (Fig. 3g,h), despite a marked reduction in RNAPII enrichment at these promoters (Extended Data Fig. 6i,j). These findings demonstrate that R-loops at the promoters of major ZGA and maternal genes form independently of RNAPII accumulation during ZGA. In summary, these R-loops are not generated through RNAPII-coupled transcription or RNAPII accumulation during ZGA.

### R-loop loss disrupts ZGA and early embryonic development

To elucidate the functional role of R-loops in the maternal-to-zygotic transition, we modulated R-loop levels by altering *Rnaseh1* expression. Microinjection of *Rnaseh1* mRNA into embryos led to global reduction of R-loops at late 2 cell stage (Fig. 4a and Extended Data Fig. 7a-d). Strikingly, R-loop loss led to embryonic developmental arrest at the 1-cell stage in a subset of embryos, accompanied by morphological degeneration (Fig. 4b-d), highlighting the essential role of R-loops in ZGA.

**Fig. 4.**
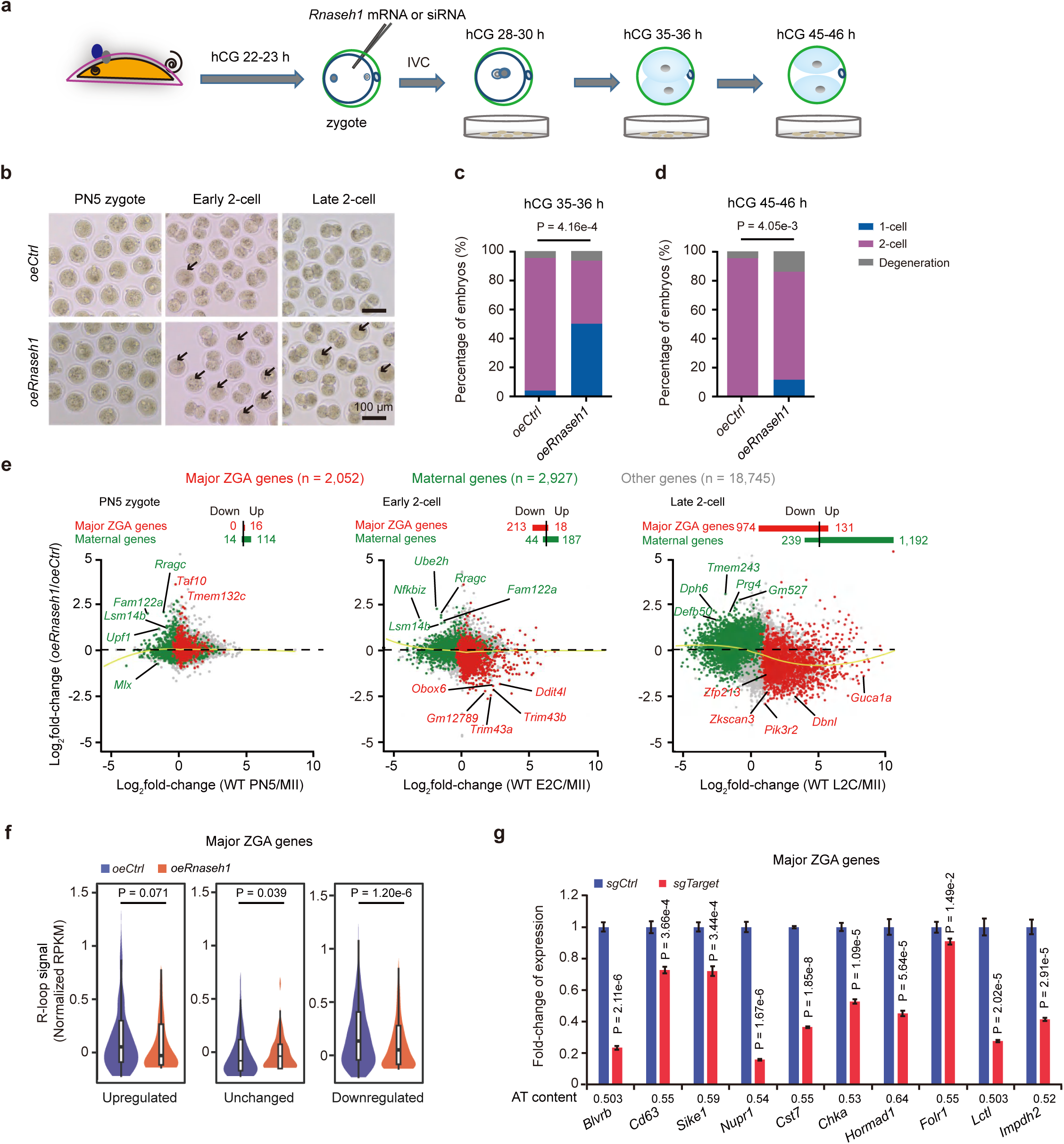
The impact of R-loops on transcriptional regulation during the maternal-to-zygotic transition. **a**, Schematic diagram illustrating the microinjection of *Rnaseh1* mRNA into early embryos followed by *in vitro* culture in KSOM medium. **b**, Representative images of embryo morphology in control (*oeCtrl*) and *Rnaseh1-*overexpressed (*oeRnaseh1*) embryos during ZGA. One representative image from three independent experiments is shown (*oeCtrl*, n = 36, 36, 30; *oeRnaseh1*, n = 54, 55, 34). Scale bars, 100 μm. Black arrows indicate abnormal embryos. **c**, Bar graphs quantifying the percentage of embryos developing to early 2-cell stage (hCG 35-36 h) with or without *Rnaseh1* overexpression (*oeCtrl*, n = 36, 36, 30; *oeRnaseh1*, n = 54, 55, 34). P value, two-sided *t*-test.| **d**, Bar graphs quantifying the percentage of embryos developing to late 2-cell stage (hCG 45-46 h) with or without *Rnaseh1* overexpression (*oeCtrl*, n = 36, 36, 30; *oeRnaseh1*, n = 54, 55, 34). P value, two-sided *t*-test. **e**, Scatter plots depicting the gene expression fold-changes upon *Rnaseh1* overexpression (two biological replicates). Yellow lines, local regression fitting. **f**, Violin plots comparing promoter R-loop enrichment (Z-score normalized) of upregulated, downregulated, and unchanged major ZGA genes with or without *Rnaseh1* overexpression. P values, two-sided Wilcoxon rank-sum test. Center line, median; box, 25th and 75th percentiles; whiskers, 1.5 × IQR. **g**, Bar graphs showing the expression levels of major ZGA gene at late 2-cell stage upon AT-rich R-loops loss with sgRNA-guided delivery of dCas9-RNase H1 to target R-loops.Corresponding AT content is indicated. P values, two-sided *t*-test. Error bars, mean ± SD.

To systematically assess the impact of R-loop perturbation on preimplantation development, we microinjected PN2 embryos with either wild-type (*WT*) *Rnaseh1* or its mutant variants, including *WKK* (mutated hybrid binding domain), *D209N* (mutated catalytical domain) and *WKKD* (mutated hybrid binding domain and catalytic domain). Microinjection of *Rnaseh1* (*WT*) caused 54.65% of embryos arrested at the 1-cell–8-cell stages, with only 3.48% developing into blastocyst stage, compared to 69.31% of blastocyst formation of embryos in control experiments (Extended Data Fig. 7e-g). Both *Rnaseh1* mutants, *WKK* and *WKKD*, had no impact on embryonic development, while *Rnaseh1* (*D209N)* mutant led to defective embryonic development (Extended Data Fig. 7e-g). The hybrid binding domain and *D209N* mutant of RNase H1 have been shown to compete with *WT* RNase H1 for R-loop binding, thereby stabilizing R-loops by preventing RNase H1 from associating with chromatin^13,23^. Thus, microinjection of *Rnaseh1* (*D209N*) also led to aberrant R-loops distribution and abnormal embryonic development, demonstrating that both R-loop loss and dysregulation disrupt early embryogenesis.

### R-loop loss leads to ectopic expression of major ZGA and maternal genes

Next, we investigated whether R-loop loss influenced the expression of major ZGA and maternal genes during ZGA. *Rnaseh1* overexpression significantly reduced genome-wide R-loop levels (Extended Data Fig. 7h,i), consistent with the immunostaining results (Extended Data Fig. 7c,d). At late 2-cell stage, *Rnaseh1* overexpression led to downregulation of 47.47% (974/2,052) major ZGA genes and upregulation of 40.72% (1,192/2,927) maternal genes (Fig. 4e). The numbers of differentially expressed genes gradually increased from PN5 zygote to late 2-cell stage (Fig. 4e), indicating that R-loop loss disrupts normal transcriptional programming during the maternal-to-zygotic transition (Extended Data Fig. 7j,k).

To further validate these findings, we next assessed the expression patterns of major ZGA and maternal genes in embryos during ZGA, where R-loops were preserved through siRNA-mediated *Rnaseh1* knockdown in PN2 embryos. While *Rnaseh1* knockdown had no significant impact on embryonic development during ZGA (Extended Data Fig. 8a-d). Transcriptome analysis revealed that *Rnaseh1* knockdown enhanced activation of major ZGA genes and accelerated silencing of maternal genes (Extended Data Fig. 8e). Notably, *Rnaseh1* knockdown promoted premature activation of major ZGA genes and facilitated the transition of maternal-to-zygotic transcription program at early 2-cell stage (Extended Data Fig. 8f-h).

Comparative analysis of differentially expressed major ZGA and maternal genes between *Rnaseh1* overexpression and knockdown at late 2-cell stage revealed reciprocal effects. Specifically, 59.72% (215/360) of the upregulated major ZGA genes in *Rnaseh1*-knockdown embryos were downregulated by *Rnaseh1* overexpression (Extended Data Fig. 8i), and 52.45% (203/387) of the downregulated maternal genes in *Rnaseh1*-knockdown embryos were upregulated by *Rnaseh1* overexpression (Extended Data Fig. 8j). Gene set enrichment analysis (GSEA) confirmed that *Rnaseh1* overexpression suppressed major ZGA gene expression while upregulated maternal genes, whereas *Rnaseh1* knockdown had the opposite effect on the expression of major ZGA and maternal genes (Extended Data Fig. 8k,l).

To explore the impact of R-loops on transcription, we analyzed the relationship between R-loop signals and gene expression, categorizing both major ZGA and maternal genes into downregulated, upregulated, and unchanged groups, respectively, based on their expression changes in late 2-cell embryos following *Rnaseh1* overexpression. Notably, in normal embryos, R-loop signals at the promoters of downregulated major ZGA genes were higher than those of the other two groups of major ZGA genes (Extended Data Fig. 8m, left). In contrast, upregulated maternal genes exhibited lower R-loop levels at their promoters compared to other maternal gene categories (Extended Data Fig. 8m, right). Moreover, *Rnaseh1* overexpression caused the most pronounced reduction of R-loops at the promoters of downregulated major ZGA genes (Fig. 4f), establishing a direct causal link between R-loop loss and transcriptional dysregulation during the maternal-to-zygotic transition.

We then investigated the specific role of AT-rich R-loops in regulating the expression of major ZGA genes during ZGA. We classified these genes into two classes based on AT content of R-loops at their promoters (AT > 0.5, AT < 0.5). Expression changes in major ZGA genes with AT-rich R-loops (AT > 0.5) showed a positive correlation with R-loop signal alterations following *Rnaseh1* overexpression (Extended Data Fig. 8n), whereas no such correlation was observed for major ZGA genes with low AT content R-loops (AT < 0.5). In contrast, maternal gene expression changes exhibited no significant correlation with R-loop alterations, regardless of R-loop AT content (Extended Data Fig. 8o).

To directly assess the functional role of AT-rich R-loops in regulating the expression of major ZGA and maternal genes, we utilized a dCas9-RNase H1 system for site-specific manipulation of AT-rich R-loops at target gene promoters. gRNA-mediated dCas9-RNase H1 targeting of selected AT-rich R-loops at major ZGA gene promoters resulted in their downregulation (Fig. 4g and Extended Data Fig. 9a,b). These findings demonstrate that R-loops, particularly AT-rich R-loops at promoters, facilitate the transition of maternal-to-zygotic transcription program, and that R-loop loss disrupts ZGA.

### R-loops promote RNAPII accumulation at major ZGA gene promoters

To elucidate the mechanism by which R-loops facilitate the maternal-to-zygotic transition, we compared R-loop levels and RNAPII signals at the promoters of unchanged and differentially expressed major ZGA genes. Downregulated major ZGA genes exhibited the highest enrichment of both R-loops and RNAPII at their promoters, in contrast to unchanged and upregulated major ZGA genes (Extended Data Fig. 8m,9d). RNAPII Stacc-seq experiments revealed that *Rnaseh1* overexpression reduced RNAPII occupancy across the promoter and gene body regions of major ZGA genes (Fig. 5a-c and Extended Data Fig. 9e). Quantitative analysis further demonstrated that the fold-change in RNAPII levels indicated a greater reduction at promoters than gene bodies of downregulated major ZGA genes (Fig. 5b,c). This was accompanied by a decreased RNAPII pausing index in *Rnaseh1*-overexpressed embryos (Extended Data Fig. 9f), suggesting that R-loop loss disrupts proper RNAPII accumulation and transcriptional regulation during ZGA.

**Fig. 5.**
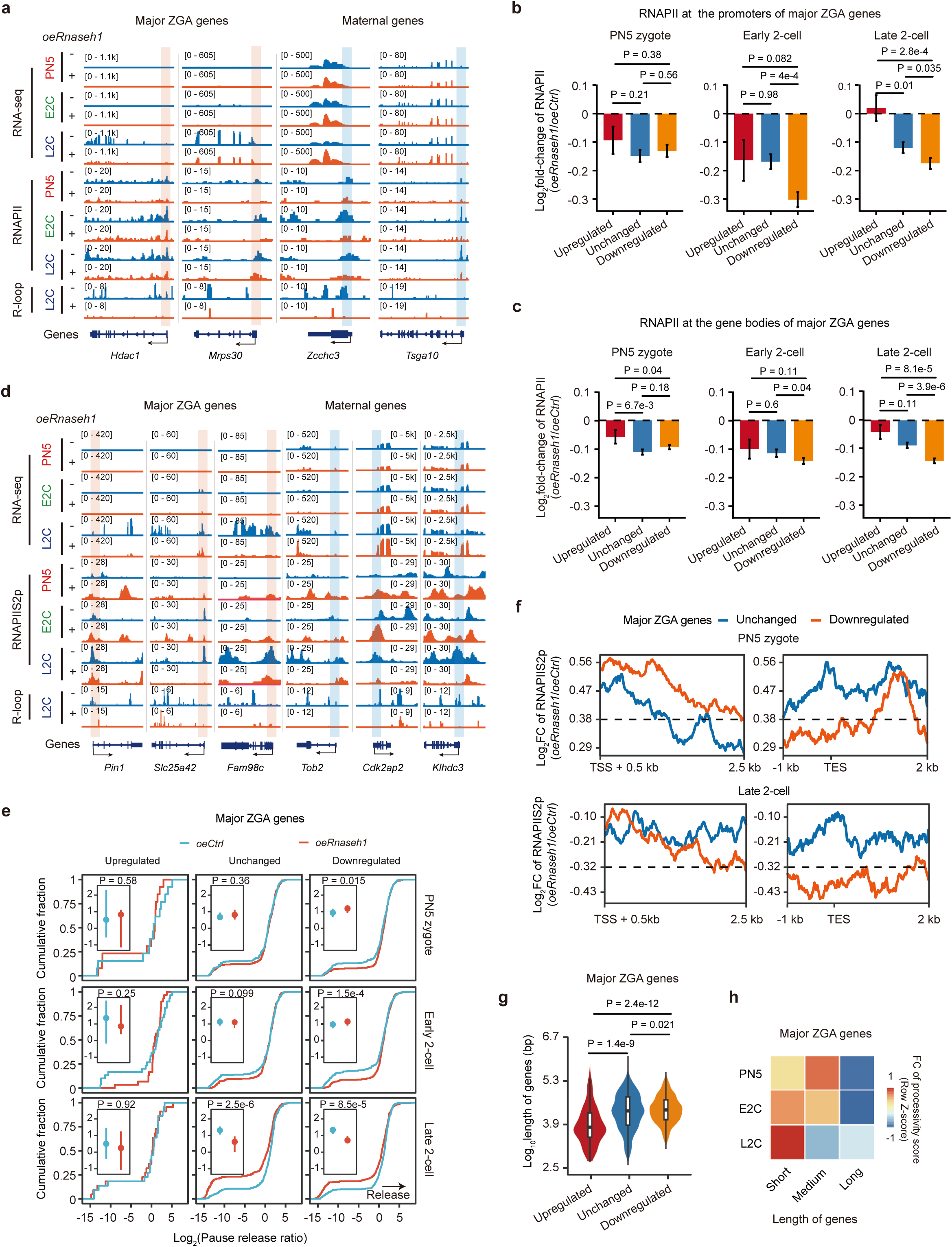
R-loops prevent pause release of RNAPII during ZGA. **a,** Tracks displaying gene expression, RNAPII occupancy, and R-loop levels in embryos with or without *Rnaseh1* overexpression during ZGA. PN5, PN5 zygote; E2C, early 2-cell; L2C, late 2-cell. **b-c**, Bar graphs showing the fold-change of RNAPII enrichment at the promoters (TSS -0.1 kb to TSS + 0.3 kb) (**b**) and the gene bodies (TSS + 0.3 kb to TES) (**c**) of upregulated, unchanged, and downregulated major ZGA genes after *Rnaseh1* overexpression during ZGA. P values, two-sided Wilcoxon rank-sum test. Error bars, mean ± SE. **d**, Tracks showing gene expression, RNAPIIS2p occupancy, and R-loop levels in embryos with or without *Rnaseh1* overexpression during ZGA. **e**, Cumulative distribution and error bar plots (insert) depicting RNAPIIS2p pause release ratios for upregulated, unchanged, and downregulated major ZGA genes in control (*oeCtrl*) and *Rnaseh1*-overexpressed (*oeRnaseh1*) embryo. P values, two-sided *t*-test. Data are presented as median ± 95% confidence intervals (CIs) in the error bar plot. **f**, Metaplots showing the fold-change (FC) of RNAPIIS2p density at 5’ regions (TSS + 0.5 kb to TSS + 2.5 kb) (left) and 3’ regions (TES -1 kb to TES + 2 kb) (right) of downregulated and unchanged major ZGA genes in control (*oeCtrl*) and *Rnaseh1*-overexpressed (*oeRnaseh1*) embryos at PN5 zygotes and late 2-cell stages. **g**, Violin plots comparing gene lengths of upregulated, unchanged, and downregulated major ZGA genes after R-loops loss through *Rnaseh1* overexpression. P values, two-sided Wilcoxon rank-sum test. Center line, median; box, 25th and 75th percentiles; whiskers, 1.5 × IQR. **h**, Heatmaps illustrating the fold-change (FC) in RNAPII processivity scores for major ZGA genes categorized by gene length (long, medium, and short) during ZGA. Data are presented as mean values.

Notably, R-loop loss resulted in a significant decrease in the RNAPII pausing index specifically at downregulated major ZGA genes at early 2-cell stage, whereas upregulated and unchanged major ZGA genes remained unaffected (Extended Data Fig. 9f). In contrast, during normal early embryonic development, the RNAPII pausing index of major ZGA genes naturally declined specifically at late 2-cell stage, coinciding with their activation at this stage (Extended Data Fig. 9g). These findings suggest that R-loop loss induces premature RNAPII pause release, potentially disrupting the precise timing of major ZGA gene activation.

The consequences of enhanced RNAPII pause release can result in either productive elongation or impaired transcriptional processivity^24,25^. Productive elongation of RNAPII typically enhances gene transcription, whereas impaired RNAPII processivity leads to transcription attenuation^25^. To distinguish between these possibilities, we measured changes in the RNAPII processivity score, defined as the ratio of RNAPII density at 3’ region of gene body (adjacent to the transcription end site, TES) to 5’ region (near the transcription start site, TSS) (Extended Data Fig. 9h). During normal early embryonic development, the RNAPII processivity score of major ZGA genes increased from early 2-cell to late 2-cell stage (Extended Data Fig. 9i). However, R-loop loss led to a reduction in the RNAPII processivity score before late 2-cell stage (Extended Data Fig. 9j). These findings suggest that R-loop loss-induced RNAPII pause release disrupts transcriptional elongation, ultimately leading to the downregulation of major ZGA genes.

### R-loop loss promotes pause release of RNAPII through aberrant phosphorylation of Ser2 in the C-terminal domain (CTD) of RNAPII

To investigate the impact of R-loop loss on RNAPII transcription activity during ZGA, we further performed Stacc-seq in *Rnaseh1*-overexpressed embryos to examine phosphorylated RNAPII at Ser2 of CTD (RNAPIIS2p), a marker of transcriptional elongation^26,27^. *Rnaseh1* overexpression resulted in a global increase in RNAPIIS2p levels at promoters and gene bodies at PN5 zygote and early 2-cell stages, followed by a subsequent decrease at late 2-cell stage (Fig. 5d and Extended Data Fig. 10a). Among unchanged and differentially expressed major ZGA genes, R-loop loss significantly elevated RNAPIIS2p occupancy at the gene bodies of downregulated major ZGA genes at PN5 zygote and early 2-cell stages but reduced RNAPIIS2p levels at late 2-cell stage (Extended Data Fig. 10b), corroborating a defect in the activation of major ZGA genes at late 2-cell stage. Notably, normally developed embryo exhibited a marked increase in RNAPIIS2p at major ZGA genes from PN5 zygote to late 2-cell stage (Extended Data Fig. 10c). In contrast, *Rnaseh1*-overexpressed embryos exhibited hyper-phosphorylation of RNAPIIS2p relative to total RNAPII at PN5 zygote and early 2-cell stages, transitioning to hypo-phosphorylation at late 2-cell stage (Extended Data Fig. 10d). The fold-change in RNAPIIS2p enrichment relative to total RNAPII at major ZGA gene bodies negatively correlated with gene expression changes (Extended Data Fig. 10e). These findings indicate that R-loop loss leads to aberrant hyper-phosphorylation of RNAPIIS2p before late 2-cell stage, ultimately impairing major ZGA activation.

To determine whether premature RNAPII pause release or hyper-phosphorylation of RNAPIIS2p before late 2-cell stage contributes to impaired RNAPII processivity on major ZGA genes, we calculated RNAPIIS2p release ratio in *Rnaseh1*-overexpressed embryos during ZGA. Downregulated major ZGA genes, but not other genes, exhibited increased RNAPIIS2p traveling downstream from TSSs at both PN5 zygote and early 2-cell stages, followed by a defect at late 2-cell stage (Fig. 5e), indicating that R-loop loss induces premature RNAPII pause release at these genes. Furthermore, *Rnaseh1* overexpression led to an elevated RNAPIIS2p signals at 5’ region of downregulated major ZGA genes at PN5 zygote stage, with a gradual attenuation toward 3’ region of genes (Fig. 5f). At late 2-cell stage, RNAPIIS2p levels in these genes were markedly reduced across the entire gene body (Fig. 5f). Collectively, these findings demonstrate that R-loop loss disrupts RNAPIIS2p dynamics, impairs elongation progression, and ultimately results in the downregulation of major ZGA genes.

RNAPII processivity is closely associated with gene length^28^. Gene length analysis revealed that downregulated major ZGA genes were significantly longer than upregulated or unchanged major ZGA genes (Fig. 5g). Notably, the RNAPII progressivity score was substantially reduced only for longer major ZGA genes following R-loop loss during ZGA (Fig. 5h). These findings suggest that R-loops preferentially facilitate the expression of longer major ZGA genes by attenuating RNAPII pause release prior to late 2-cell stage and ensuring processive elongation.

We next investigated whether the resolution of a single R-loop at promoter could alter RNAPIIS2p levels on major ZGA genes in early embryos. Utilizing site-specific R-loop editing with dCas9-RNase H1, we found that the removal of a single R-loop led to increased RNAPIIS2p enrichment at both promoters and gene bodies of major ZGA genes. Notably, the enrichment was more pronounced at promoters and 5’ regions of genes compared to 3’ regions (Extended Data Fig. 10f). The failure of RNAPIIS2p to efficiently travel to transcription end sites (TESs) correlated with the downregulation of major ZGA genes (Fig. 4h and Extended Data Fig. 10f). These results demonstrate that R-loop loss disrupts RNAPII pause release and processive activity.

### R-loops attenuate DDX21 helicase activity on the 7SK/HEXIM1 snRNP complex

To elucidate the mechanism in which R-loops regulate RNAPII pause release during ZGA, we analyzed mass spectrometric data of RNAPIIS2p and RNAPIIS5p interactomes in mouse embryonic stem (mES) cells. DDX21, a helicase known for resolving R-loops^29^, promoting RNAPII pause release^30,31^ and facilitating transcription termination^32^, was significantly enriched in both RNAPIIS2p and RNAPIIS5p interactomes (Extended Data Fig. 11a). At promoters, DDX21 facilitates the release of positive transcription elongation factor b (P-TEFb) from the 7SK/HEXIM1 snRNP complex through its helicase activity, thereby transitioning CDK9 from inactive to active^30^.

During normal embryonic development, DDX21 enrichment increased at major ZGA gene promoters (Extended Data Fig. 11b). Additionally, DDX21 binding to 7SK snRNA exhibited an increasing trend during ZGA (Extended Data Fig. 11c). Meanwhile, HEXIM1, inhibiting RNAPlI elongation by 7SK/HEXIM1 snRNP complex formation^33^, showed increased binding at major ZGA gene promoters from PN5 zygote to early 2-cell stage, followed by decreased binding at late 2-cell stage (Extended Data Fig. 11d). These findings suggest that DDX21 and HEXIM1 exhibit distinct binding patterns at major ZGA genes.

We next investigated whether DDX21 is required for R-loop-mediated regulation of promoter-proximal RNAPII pause release. While R-loop loss induced by *Rnaseh1* overexpression did not affect DDX21 binding at major ZGA gene promoters at PN5 zygote stage (Fig. 6a), it significantly reduced DDX21 enrichment at late 2-cell stage (Extended Data Fig. 11e). These findings suggest that R-loop-dependent recruitment of DDX21 at the promoters of major ZGA genes.

**Fig. 6.**
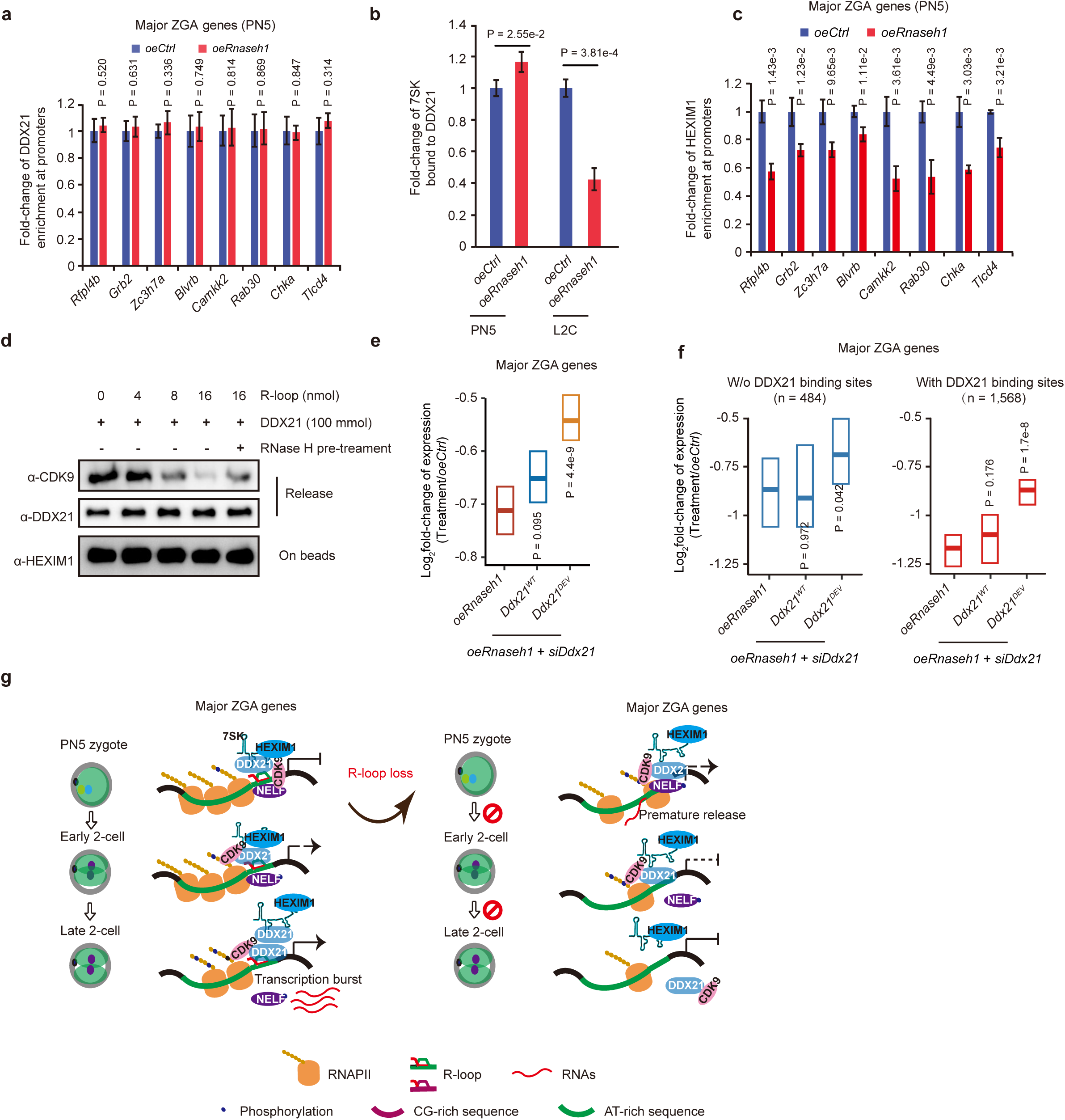
R-loops attenuate DDX21 helicase activity on the 7SK/HEXIM snRNP complex to promote expression of major ZGA genes and inhibit expression of maternal genes. **a**, Bar graphs depicting promoter DDX21 enrichment of major ZGA genes in control (*oeCtrl*) and *Rnaseh1*-overexpressed (*oeRnaseh1*) embryos at PN5 zygote (PN5) stage.P values, two-sided *t*-test. Error bars, mean ± SD. **b**, Bar graphs illustrating the enrichment of 7SK bound to DDX21 in control (*oeCtrl*) and *Rnaseh1*-overexpressed (*oeRnaseh1*) embryos at PN5 zygote (PN5) and late 2-cell (L2C) stages. P values, two-sided *t*-test. Error bars, mean ± SD. **c**, Bar graphs depicting promoter HEXIM1 enrichment of major ZGA genes in control (*oeCtrl*) and *Rnaseh1*-overexpressed (*oeRnaseh1*) embryos at PN5 zygote (PN5) stage.P values, two-sided *t*-test. Error bars, mean ± SD. **d**, Western blot analysis of a P-TEFb release assay (measured as CDK9 levels) with DDX21 and varying amounts of synthetic R-loop oligos. **e**, Crossbar plots showing the expression fold-change of major ZGA gene in embryos with *Rnaseh1* overexpression (*oeRnaseh1*), *Rnaseh1* overexpression combined with endogenous *Ddx21* knockdown rescued by either wild-type *Ddx21* (*Ddx21^WT^*) or a helicase activity defective mutant *Ddx21^DEV^*, relative to the control. Each group is compared to *oeRnaseh1*. P values,two-sided Wilcoxon rank-sum test. Data are presented as median ± 95% CIs. **f**, Crossbar plots displaying the expression fold-change of major ZGA genes with or without DDX21 binding sites following *Rnaseh1* overexpression (*oeRnaseh1*), *Rnaseh1* overexpression combined with endogenous *Ddx21* knockdown (*siDdx21*) rescued by either wild-type *Ddx21* (*Ddx21^WT^*) or a helicase activity defective mutant *Ddx21^DEV^*, relative to the control. Each group is compared to *oeRnaseh1*. P values, two-sided Wilcoxon rank-sum test. Data are presented as median ± 95% CIs. **g,** A model illustrating the role of R-loops in gene expression and early embryonic development. At the pre-ZGA stage, R-loops inhibit RNAPII pausing release at maternal genes, contributing to the silencing of these genes. Conversely, at major ZGA genes, R-loops accumulate RNAPII by inhibiting DDX21 helicase activity on the 7SK/HEXIM1 snRNP complex, maintaining CDK9 in its inactive state at PN5 zygote and early 2-cell stages. Upon R-loop resolution by DDX21, enhanced binding of DDX21 to 7SK/HEXIM1 promotes CDK9 release, leading to major ZGA genes activation at late 2-cell stage. R-loop loss leads to abnormal early embryonic development and aberrant gene expression during the maternal-to-zygotic transition.

We further examined the impact of R-loop loss on the association between 7SK snRNA and DDX21, as well as HEXIM1 occupancy at the promoters of major ZGA genes. R-loop loss significantly increased the interaction between 7SK snRNA and DDX21 at PN5 zygote stage but led to a decrease at late 2-cell stage (Fig. 6b). Additionally, HEXIM1 enrichment at the promoters of major ZGA genes was reduced at PN5 zygote and late 2-cell stages following R-loop loss (Fig. 6c and Extended Data Fig. 11f), aligning with the enhancement of RNAPII pause release.

Given that DDX21 facilitates RNAPII pause release in a helicase activity-dependent manner^30^, we hypothesized that R-loops might compete with 7SK snRNA for DDX21 binding, thereby attenuating helicase activity of DDX21 on the 7SK/HEXIM1 snRNP complex. To test this, we purified 7SK/HEXIM1 snRNP complex from mES cells and incubated it with DDX21 along with varying amount of synthetic R-loop oligonucleotides. Our results showed that R-loops inhibited CDK9 release from the 7SK/HEXIM1 snRNP complex in a dose-dependent manner, whereas RNase H pre-treatment enhanced CDK9 release by DDX21 (Fig. 6d). These findings demonstrate that R-loops suppress DDX21 helicase activity on the 7SK/HEXIM1 snRNP complex, thereby maintaining CDK9 in inactive state and promoting RNAPII pausing at promoters.

### Loss of DDX21 helicase activity rescues gene expression alterations induced by R-loop loss

To further elucidate the functional role of DDX21 helicase activity in R-loop-mediated gene regulation, we generated a helicase defective DDX21 mutant (DDX21^DEV^, which lacks catalytic activity)^34^ and assessed its impact on major ZGA and maternal gene expression in *Rnaseh1*-overexpressed embryos following endogenous *Ddx21* knockdown. Notably, expression of major ZGA genes downregulated by R-loop loss was effectively restored by DDX21^DEV^ (Fig. 6e, Extended Data Fig. 11g) accompanied by the rescued level of maternal transcripts (Extended Data Fig. 11h). These findings demonstrate that DDX21 helicase activity is essential for R-loop-mediated transcriptional regulation of both major ZGA and maternal genes.

We then examined the impact of DDX21 binding on the expression of major ZGA l genes. *Rnaseh1* overexpression led to a more pronounced decrease in R-loop levels at the promoters of major ZGA genes containing the DDX21 binding compared to those lacking DDX21 binding (Extended Data Fig. 12a). Major ZGA genes with promoters harboring the DDX21 binding exhibited greater expression downregulation following R-loop loss (Fig. 6f). Notably, major ZGA genes containing the DDX21 binding showed higher rescue potential (Fig. 6f), suggesting that DDX21 helicase activity is essential for R-loop dependent activation of major ZGA genes. Furthermore, in *Rnaseh1*-overexpressed embryos, the transcriptional rescue efficiency by DDX21^DEV^ in major ZGA genes harboring promoter AT-rich R-loops negatively correlated with R-loop changed levels (Extended Data Fig. 12b). Moreover, maternal transcripts were rescued in *Rnaseh1*-overexpressed late 2-cell embryos upon depletion of endogenous *Ddx21* combined with *Ddx21^DEV^* overexpression (Extended Data Fig. 12c).

In conclusion, R-loops, particularly AT-rich R-loops, act as key regulators of RNAPII pausing by inhibiting DDX21 helicase activity. This mechanism ensures proper silencing of maternal transcription program by preventing premature RNAPII pause release at major ZGA genes, thereby facilitating the transcriptional burst for zygotic transcription program (Fig. 6g).

## Discussion

Previous studies have reported that R-loops serve as multifaceted regulators in cell fate determination^13,35,36^ and RNase H deficiency is known to result in embryonic lethality^12^. However, the reprogramming dynamics of R-loops during mammalian preimplantation embryonic development remains enigmatic due to technical limitations in profiling low-input embryonic material. Our study provides comprehensive atlas of R-loop dynamics in mouse gametes and preimplantation embryos, uncovering fundamental differences from mammalian cell lines^16^. Notably, we identified predominant AT-rich R-loops during early embryogenesis, exhibiting stage-specificity and regulatory potential. Rather than being mere transcriptional byproducts, these R-loops function as transient epigenetic elements, highlighting their critical role in early developmental processes.

Our findings reveal a dual regulatory mechanism in which R-loops orchestrate transcriptional priming at major ZGA genes while simultaneously enforcing maternal gene silencing. Notably, we observed a slight decrease in R-loop levels following the abundant accumulation of RNAPII or transcription activation (Fig. 3d), suggesting a self-limiting regulatory circuit, potentially mediated by transcription-coupled recruitment of R-loop resolution factors^23,37^. This dynamic equilibrium highlights the paradoxical nature of R-loops: while essential for transcriptional regulation, their persistence necessitates strict control through dedicated surveillance mechanisms^1,2^. The differential sensitivity of major ZGA and maternal genes to these regulatory pathways likely reflects their distinct R-loop burdens and promoter architectures. Moreover, the unexpected regulatory role of R-loops in a subset of maternal gene expression during ZGA raises important questions about the dynamics of maternal genes during the maternal-to-zygotic transition. This is consistent with previous studies showing that the re-expression or replenishment of maternal transcripts also occurs during this critical developmental period^38,39^.

The RNAPII-uncoupled nature of R-loop formation, as evidenced by DRB and α-amanitin treatments, suggests the involvement of alternative RNA sources. Several potential mechanisms may account for this phenomenon: 1) DRB or α-amanitin inhibits RNAPII transcription at highly activated genes in wild-type embryos; however, the suppression of RNAPII activity induce promiscuous ectopic transcription at other genomic regions^22^. The resulting ectopic RNAs could hybridize with DNA to form R-loops. 2) Transcription is not entirely abolished following DRB/α-amanitin treatment, as residual transcriptional activity from RNA polymerase I/III may persist. Notably, RNA polymerase I/III have been reported to transcribe certain cellular mRNAs^40,41^, as well as small RNAs at protein-coding gene promoters^42^, potentially contributing to R-loop formation. 3) Maternal RNAs might hybrid with genomic DNA during DNA replication or the DNA double-strand break repair process. Additionally, it is undeniable that this study, like others investigating early mouse embryos, may miss a fraction of low-abundance R-loops due to the bias of RIAN-seq, which preferentially captures high-signal R-loops in low-input cell samples. Similar limitations have been reported for other methods, such as Stacc-seq^22^, STAR-ChIP^43^, mini-ATAC-seq^44^.

Taken together, our study establishes R-loops as critical temporal regulators of early mouse embryonic development. In particular, AT-rich R-loops play a dual role by facilitating the silencing of maternal transcriptional program while preserving transcriptional competence at major ZGA genes. This coordinated regulation ensures the fidelity of maternal-to-zygotic transition, effectively balancing the paradoxical demands of transcriptional quiescence and rapid activation during early development. Our findings not only deepen our understanding of R-loops as potential biomarkers for embryo viability but also opens new avenues for investigating their regulatory roles in other developmental processes and disease contexts.

## Methods

No statistical methods were used to predetermine sample size. The experiments were not randomized, and the investigators were not blinded to allocation during the experimental procedures and outcome assessment.

### Collections of mouse gametes and early embryos

Early embryos were collected from C57BL/6 female mice mated with PWK/PhJ male mice. 5-8 weeks C57BL/6 females were induced to superovulation by intraperitoneal injection of hCG (10 IU) at an interval of 46-48 h after PMSG (10 IU) injection. Oocytes and embryos were collected at the following time points after hCG injection: MII oocytes (13 h, no mating), PN2 zygote (21-23 h), PN5 zygote (28-30 h), early 2-cell (35-36 h), late 2-cell (45-46 h), 4-cell (54-56 h), 8-cell (66-68 h), and blastocysts (98-100 h). The inner cell mass (ICM) was then separated from blastocysts. Granulosa cells were removed by treatment with 0.5% hyaluronidase, while the zona pellucida was removed with Tyrode’s solution (Sigma, T1788). Sperm collection experiment was performed according to a previous study^45^ with minor modifications. Briefly, sperm was obtained from 12-week-old PWK/PhJ males by the swim-up and treated with somatic lysis buffer. DNase I (NEB, M0303S) was used to remove cell-free DNA. The purified sperm was then treated with DTT. All procedures using animals were reviewed and approved by the Institutional Animal Care and Use Committee (IACUC) of Guangzhou National Laboratory. All experimental procedures were performed in accordance with the Principles for the Care and Use of Laboratory Animals.

### DRB and α-amanitin treatment of mouse embryos

PN2 embryos were obtained 23 h after hCG injection and then cultured in KSOM medium supplemented with 100 μM α-amanitin or 60 μM DRB. At the same time, the control group was treated with ddH_2_O or 0.1% DMSO, respectively. These embryos were collected at 14 h (early 2-cell) or 25 h (late 2-cell) after treatment. All *in vitro* culture experiments were carried out under 5% CO_2_ in a 37°C incubator.

### Microinjection of *Rnaseh1/Ddx21* mRNA and siRNA into early embryos

For mRNA, the coding sequence (CDS) of *Rnaseh1* (*WT*) and *Rnaseh1* (*WKK/D209N/WKKD*) mutants, *Ddx21* (*WT*) and *Ddx21* (*DEV*) mutants, and *dCas9-Rnaseh1* were cloned into the pGEMHE vector with a T7 promoter. After linearization, the plasmid fragments were retrieved. mRNA was transcribed using mESSAGE mMACHINE T7 Kit (Invitrogen, AM1344) or T7 High Yield RNA Transcription Kit (Vazyme, TR101) with the CAG Trimer Cap analogue. For sgRNA, the upstream 20 bp of the NGG protospacer adjacent motif (PAM) sequences used as the sgRNA sequences were recombined into the pUC57kan-T7-gRNA vector. After amplification, the sequences were transcribed by T7 High Yield RNA Transcription Kit (Vazyme, TR101). All micromanipulations were performed using a Piezo impact-driven micromanipulator. PN2 zygotes were temporarily cultured in M2 medium and 5-10 pL 400 ng/μL *Rnaseh1* (WT and mutants) mRNA or 12.5 μM siRNA was microinjected per embryo. For the DDX21 rescue experiment, 12.5 μM *Ddx21* siRNA and 200 ng/μL *Ddx21* (*WT*) or *Ddx21* (*DEV*) were microinjected after microinjecting *Rnaseh1*. For site-specific regulation of R-loops, 400 ng/μL *dCas9-Rnaseh1* and 60 ng/μL sgRNA were microinjected, respectively. Then the embryos were transferred into KSOM medium and cultured until collection for further experiments. The sequences of siRNA were shown in Supplementary Table 2.

### RIAN-seq

Genomic DNA was purified from cells or embryos using a genomic DNA extraction kit and then incubated with NEB restriction enzyme mix (Alu I, Mbo I, Mse I, Dde I), Nuclease P1, T5 exonuclease, and Lambda exonuclease at 37°C for 1 h. As controls, RNase H (NEB, M0297S) was added during the incubation stage. 0.2 µL of Proteinase K (Thermo Scientific, EO0491) was added before incubation at 55°C for 30 min and 70°C for 30 min. 0.3 µL 1 M Tris-HCl (pH 8.0), 0.15 µL 1M MgCl_2_, 1.5 µL PEG200, 4.8 µL 50% PEG8000 and 0.5 µL transposase (Vazyme, TD503) were added and incubated at 55°C for 10 min. The transposition was stopped by the addition of 5 µL 0.25% SDS and incubation at room temperature (RT) for 5 min. 1 µL10% Triton X-100 was added and incubated at 37°C for 1 h to quench SDS. Transposed DNA was gap-filled by addition of 2 µL *Bst* 3.0 DNA polymerase (NEB, M0374S) and 25 µL NEBNext Q5 Hot Start HiFi PCR Master Mix (NEB, M0543S). The mixture was incubated at 72°C for 15 min and 80°C for 10 min. For library amplification, the mixture was mixed with a barcoded i5 primer and a barcoded i7 primer, and PCR was performed to amplify the libraries using the following conditions: 98°C for 45 s; thermocycling for 10 cycles at 98°C for 15 s, 65°C for 75 s; followed by 65°C for 5 min. The final libraries were purified with the 1 × AMPure XP beads (Beckman, A63882) size selection and were subjected to next-generation sequencing.

### Stacc-seq

Stacc-seq experiments were performed according to previous study^22^ with a few modifications. Briefly, the embryos were resuspended in 50 μL PBS containing 0.005% Digitonin. After incubation at 4°C for 10 min, 8 μL pre-assembled antibody-pG-Tn5 complex (containing 5 μL PBS, 2.5 μL RNAPII (or RNAPIIS2p) antibody, 0.5 μL pG-Tn5 or pA-Tn5) and 14.5 μL 5 × TTBL (Vazyme, TD501) were added. The samples were incubated at 37°C for 30 min, and 2 μL 0.5 M EDTA, 0.5 μL 20% SDS and 2 μL 20 mg/mL proteinase K were added. Then the tubes were incubated at 55°C for 15 min. After extraction with phenol-chloroform followed by ethanol precipitation, the samples were prepared for PCR. Library generation was performed under the following the conditions: 72°C for 5 min; 98°C for 45 s; and thermocycling for 16 cycles at 98°C for 15 s, 60°C for 30 s and 72°C for 3 min; followed by 72°C for 5 min. After the PCR reaction, libraries were purified with AMPure XP beads (Beckman).

### RNA-seq

The RNA-seq libraries were generated from embryos using Geo-seq^46^ as described previously with minor modifications. Embryos were lysed in GuSCN (Guanidine isothiocyanate) solution, and the polyadenylated mRNAs were captured with the PolyT primers. Then RNA samples were incubated for denaturation at 72°C for 3 min and used for reverse transcription. After pre-amplification and purification with AMPure XP beads, libraries were generated using the TruePrep DNA Library Prep Kit V2 for Illumina (Vazyme, TD503) according to the manufacturer’s instruction. All libraries were sequenced on NovaSeq 6000 according to the manufacturer’s instructions.

### Immunostaining

Zona pellucida was removed from oocytes or embryos by using Tyrode’s solution. Oocytes and embryos were fixed with 4% paraformaldehyde (PFA) at RT for 30 min. After 3 times washing with PBS-T (0.1% Triton X-100 in PBS), oocytes and embryos were permeabilized with 0.5% Triton X-100 in PBS at RT for 1 h. For detection of RNase H1 signal, oocytes and embryos were incubated with 3% BSA at RT for 1 h. Primary antibodies were incubated overnight at 4°C (RNase H1 antibody, Proteintech, 15606-1-AP), and then the oocytes or embryos were washed 3 times with PBS-T (0.1% Triton X-100 in PBS). For detection of R-loop with S9.6 antibody, oocytes and embryos were treated with 4 M HCl at RT for 10 min and then neutralized with 150 mM Tris-HCl (pH 8.0). For RNase H treatment, oocytes and embryos were treated with 200 U/mL RNase H (NEB, M0297S) for 2 h at 37°C. After three washes with PBS-T (0.1% Triton X-100 in PBS), the oocytes and embryos were blocked with 3% BSA at RT for 1 h. Primary antibodies were incubated overnight at 4°C (S9.6 antibody, homemade), and then the oocytes or embryos were washed 3 times with PBS-T (0.1% Triton X-100 in PBS) on a shaker. The secondary antibody was incubated at RT for 1 h. After 3 times washing with PBS-T (0.1% Triton X-100 in PBS), the samples were incubated with DAPI or PI to label the nucleus. LSM 800 or 900 (ZEISS) was used for image collection, and images were analyzed with ImageJ or ZEN software. The primary antibodies are listed in Supplementary Table S3. The secondary antibodies used in this study are as follows: Goat anti-Mouse Alexa Fluor 488 (Invitrogen, A11001), Goat anti-Rabbit Alexa Fluor 488 (Invitrogen, A11008), and Goat anti-mouse Alexa Fluor 647 (Invitrogen, A21235).

### Quantitative analysis of RNAPII, RNAPIIS2p, DDX21, HEXIM1 enrichment at promoters and 7SK bound to DDX21

Stacc-seq and CUT&RUN were performed according to previous reports^22,47^. 2 μL Spike-in DNA was mixed with DNA from Stacc-seq or CUT&RUN. qPCR primers used in this study were listed in Supplementary Table 2.

### RIAN-seq data processing

Adapters and low-quality bases of raw reads were trimmed using TrimGalore (v0.6.1). Then, the reads were aligned to mm10 reference genome using Bowtie2 (v2.5.1) with the parameters: -N 1 -L 25 -X 2000 -p 16 --no-mixed --no-discordant. Low mapping quality reads (MAPQ < 20) and PCR duplicates were removed using SAMtools (v1.16.1)^48^ and Picard (v2.2.4) (https://broadinstitute.github.io/picard/). Only the uniquely mapped reads without PCR duplicates were retained. Coverage tracks were generated using BEDTools (v2.29.2)^49^ and bedGraphToBigWig (http://hgdownload.cse.ucsc.edu/admin/exe/). To enhance signal quality, coverage tracks were further denoised with AtacWorks (v0.3.0)^15^, a deep learning model trained on high-quality RIAN-seq data to reduce background noise. Reads per kilobase of bin per million reads sequenced (RPKM) value was calculated in 100-bp bins. To reduce the batch effects and cell type variation, RPKM values were further normalized through Z-score transformation. The *Z* score was obtained using the following formula: for a given bin *i*: *z_i_* = (*x_i_* − *μ*)/*σ*, where *x_i_* represents the RPKM value before normalization, *z_i_* is the normalized RPKM value and *μ* and *σ* are the mean and standard deviation of all RPKM values (excluding blacklisted regions) for each stage, respectively. The correlation between biological replicates was calculated using multiBamSummary and plotCorrelation functions from deepTools (v3.4.3)^50^.

### Model evaluation in AtacWorks

To evaluate the denoising performance of AtacWorks (v0.3.0)^15^, BAM files for bulk RIAN-seq derived from HEK293T cells were downsampled to a fixed read count using SAMtools (v1.16.1). Low coverage data were denoised using Atacworks (v0.3.0). The Pearson correlation coefficient between bulk RIAN-seq and subsampled datasets was computed before and after denoised with Atacworks (v0.3.0). To evaluate peak-calling performance on low-coverage data, the Area Under the Receiver-Operator Characteristic (AUROC) was calculated for MACS2 (v2.1.0) and AtacWorks (v0.3.0).

### Hierarchical clustering analysis

Hierarchical clustering was performed using the hclust() function in R (v4.2.3).

### Stacc-seq data processing

Stacc-seq data were processed and peaks were called as previously described^22^. To minimize the batch effects and cell type variation, RNAPII signals were quantified as RPKM values and further normalized by the Z-score transformation.

### Rank normalization of data

To better quantify differences in RNAPII or R-loop occupancy between *Rnaseh1*-overexpressed and control samples, a rank normalization (quantile normalization)^51^ was applied to normalize the datasets. Briefly, for each dataset, genomic bin (100-bp) were ranked based on signal intensity (RPKM). The highest-ranked bin in each dataset was assigned the average signal of all top-ranked bins across datasets. This process was iterated across all bins.

### RNA-seq data processing

Adapter sequences and low-quality bases in raw reads were trimmed using TrimGalore (v0.6.1) with parameters: --stringency 10 --max_n 10 --length 25 –paired. Processed reads were then mapped to mm10 genome reference using HISAT2 (v2.1.4)^52^, with options --dta-cufflink. Differential gene expression analysis between treatment and controls was performed with Cuffdiff function of Cufflinks (v2.2.1) with parameters: -u --min-reps-for-js-test 2. Genes with q value < 0.05 and absolute log_2_fold-change > 0.58 were classified as differentially expressed genes (DEGs).

### Identification of promoter and distal R-loops

R-loop peaks were identified usinghe probability track generated by AtacWorks (v0.3.0), applying a cutoff of 0.9. Peaks were called using *peaksummary.py* script provided by AtacWorks (v0.3.0). Only peaks with strong signals (coverage > 15) and those not overlapping with a custom-build blacklist were retained for further analysis. Peaks located within ± 2.5 kb of the TSS range were defined as promoter R-loop peaks, while those positioned at least 3.5 kb away from TSSs were defined as distal R-loop peaks.

### Distribution of R-loop peaks

To compare the enrichment of R-loop peaks on different genomic elements, promoters (TSS ± 2.5 kb), exons, introns, terminator regions (TES ± 2.5 kb) and intergenic regions were defined using Ensembl annotation. R-loop peaks were assigned to each group based on the center of the peaks with the following priority: promoter, exon, intron, terminator regions, intergenic.

### Identification of stage-specific R-loops

The identification of stage-specific R-loops was performed as previously reported^44^ with slight modifications. Briefly, R-loop peaks from gamete to ICM were merged, and average Z-score normalized RPKM values were assigned. Stage-specific R-loop candidates were then identified using a Shannon entropy-based method, selecting peaks with an entropy value of less than 1. Stage-specific R-loops were defined if the candidates met the following criteria: The R-loop peaks exhibited high enrichment at a specific stage (normalized RPKM > 2), while no comparable signal (normalized RPKM > 2) was detected at other developmental stages.

### Definition of the gained, lost and shared R-loops

To analyze the dynamic changes in R-loops, peaks from adjacent developmental stages were merged, and average Z-score normalized RPKM values were computed. Peaks exhibiting high R-loop enrichment at the current stage (normalized RPKM > 2) but low enrichment at the previous stage (normalized RPKM < 2) were defined as gained R-loops. In contrast, peaks with high enrichment at the previous stage (normalized RPKM > 2) but low enrichment at the current stage (normalized RPKM < 2) were defined as lost R-loops. Shared R-loops were identified when the normalized RPKM exceeded 2 at both stages.

### Comparison between R-loops and other epigenetic modifications

The raw FASTQ files of RNAPII, H3K36me3, H3K9me3, ATAC-seq, H3K4me3 and H3K27me3 were downloaded from the GEO database (accession numbers: GSE135457, GSE112834, GSE98149, GSE66581, GSE71434 and GSE76687, respectively). These data were aligned to the mouse genome (mm10) using Bowtie2 (v 2.5.1). Uniquely mapped reads without PCR duplicates were retained and used to generate bigWig files containing Z-score normalized RPKM. The level of colocalization was measured by calculating the enrichment of RNAPII, H3K36me3, H3K9me3, ATAC-seq, H3K4me3 and H3K27me3 at promoter and distal R-loops using the multiBigwigSummary function of deepTools (v3.4.3).

### Definition of high, intermediate and low CpG promoters

Promoters were classified as previously described^53,54^. The CpG ratio was first calculated for 500-bp bins with 50-bp steps using the formula: (number of CpGs * number of bp) / (number of Cs * number of Gs). Based on the CpG ratio and GC content cutoff, promoters (within ± 2.5 kb of the TSS) were classified into three categories: high CpG promoters (HCP) (containing a 500-bp interval with a CpG ratio greater than 0.6 and GC content higher than 55%), low CpG promoters (LCP) (not containing a 500-bp interval with a CpG ratio greater than 0.4), and intermediate CpG promoters (ICP) (neither HCP nor LCP). Average R-loop signals (normalized RPKM) were quantified across all promoter types and visually represented.

### Identification of maternal genes, major ZGA genes and PcG target genes

The RNA-seq data of early mouse embryos were downloaded from the GEO database (GSE71434). Major ZGA genes were defined as those significantly upregulated in late 2-cell embryos (FPKM > 1 and fold-change > 3 compared to PN5 zygote), based on a previous report^55^. To avoid potential confounding effects from maternally inherited RNA transcripts, only genes not expressed in MII oocytes (FPKM < 0.5) were retained for downstream analysis.

Genes with high expression levels (FPKM > 5) at MII oocyte stage were retained for identifying maternal genes. The expression level of each gene was transformed by log_2_(FPKM + 1). Maternal genes were identified based on the following criteria^56^: (1) expression (MII) > expression (L2C) + 1, expression (L2C) < expression (4-cell) + 1, expression (L2C) > expression (4-cell) − 1; (2) expression (MII) < expression (L2C) + 1, expression (MII) > expression (L2C) − 1, expression (L2C) > expression (4-cell) + 1; (3) expression (MII) > expression (L2C) + 1, expression (L2C) > expression (4-cell) + 1. Genes marked by H3K27me3 at their promoters (TSS ± 2.5 kb) in mES cells were classified as PcG target genes^57^. Genes with FPKM > 3 or < 0.5 in all stages were considered constantly active or inactive genes, respectively.

### Comparison between RNAPII and R-loop enrichment at promoters with different AT contents

The AT content of promoters was profiled using nucBed in BEDTools (v2.29.2). Promoters were then divided into five groups based on their AT content (0.40-0.45, 0.45-0.50, 0.50-0.55, 0.55-0.60, 0.60-0.65). These groups collectively accounted for 97.8% of all genomic promoters. The Pearson correlation between RNAPII and R-loop enrichment (Z-score normalized) across the different promoter groups was computed and visualized in a line chart.

### Calculation of pause release ratio (PRR) and pausing index

The promoter region (100 bp upstream of the TSS to 300 bp downstream of the TSS) and gene body region (300 bp downstream of the TSS to 2 kb downstream of the TSS) used for computing PRR and pausing index were defined as previously reported^58^. The PRR was calculated as the level of RNAPII within the gene body divided by the level of RNAPII within the promoter. The pausing index was calculated as the RNAPII level within the promoter divided by the RNAPII level within the gene body.

### Calculation of transcription processivity score

The processivity score of a given gene was computed as previously reported^59^. The processivity score was defined as the log_2_fold-change of the RNAPII density in the distal region (1 kb upstream of the TES to 2 kb downstream of the TES) versus the proximal region (500 bp downstream of the TSS to 1.5 kb downstream of the TSS).

### Estimation of DNA methylation levels of R-loops

The DNA methylation data were downloaded from the GEO database (GSE98151 and GSE56697). The DNA methylation level for each R-loop peak was quantified as the average DNA methylation level of all CpG sites covered within the region. The DNA methylation level of each gene promoter region was computed in the same manner, but only promoters with at least 5 CpG sites covered were included in the analysis.

### DDX21 binding site prediction

DDX21 binding sites around TSSs (TSS ± 2.5 kb) of major ZGA and maternal genes were predicted using PrismNet^60^, a deep learning model based on RNA sequence and *in vivo* RNA structure data. PrismNet was trained on the eCLIP data of DDX21 from ENCODE^61^, along with the matched RNA structural data (icSHAPE scores) in K562 (https://zhanglabnet.oss-cn-beijing.aliyuncs.com/prismnet/correlation_shape_rpkm/K562_smartSHAPE.out). For input, the sequences and the icSHAPE data of mESCs (https://zhanglabnet.oss-cn-beijing.aliyuncs.com/prismnet/correlation_shape_rpkm/mES_smartSHAPE.out) were split into sliding windows (window size: 101 nt, step: 20 nt). The sequence window with a binding probability over 0.6 was regarded as a predicted binding site of DDX21. To better analyze the relationship between R-loops and DDX21, only the predicted binding sites with high R-loop enrichment (normalized RPKM >1) were retained. Overlapping binding sites were merged for further analysis.

### Gene set enrichment analysis (GSEA)

The GSEA analysis was performed on the gene sets of major ZGA and maternal genes using clusterProfiler R package (v4.14.4)^62^.

### GO analysis

Gene Ontology enrichment analysis for genes near stage-specific R-loop peaks was performed using Genomic Regions Enrichment of Annotations Tool (GREAT) (http://great.stanford.edu/public/html/).

## Data availability

All data have been deposited in the Genome Sequence Archive under project XXXX, with the accession number XXXX. The sample information, including sample name, developmental stage, cell number, treatment, and experiment, is summarized in Supplementary Table 1.

## Code availability

Software and code used to analyze these data are listed in the Nature Research Reporting Summary and are all publicly available.

## Acknowledgements

This work was supported by the National Key R&D Program of China (2021YFA1100300), the National Natural Science Foundation of China (32430016, 32200456, U21A20195), Major Project of Guangzhou National Laboratory (GZNL2023A02010, GZNL2023A02008), and the Science and Technology Planning Project of Guangdong Province, China (2021A1515110181, 2021A1515111061).

## Author contributions

H.Y. and Y.L. conceived and designed the experiments. Y.L. performed RIAN-seq and Stacc-seq experiments. Q.L. and Y.S. conducted bioinformatics analysis. X.W., C.D. and S.H. collected gametes and early mouse embryos, and performed RNA-seq experiments. X.W. conducted microinjection and immunostaining with the help of Q.C. C.D. constructed the plasmid construction. J.C., G.W., S.G. provided mice and technical supports. H.Y., Y.L., Q.L. and X.W. wrote the manuscript. H.Y. funded and supervised the entire study.

**Supplementary Information** is available for this paper.

**Correspondence and requests for materials** should be addressed to H.Y.

---

**Extended Data Fig. 1.**
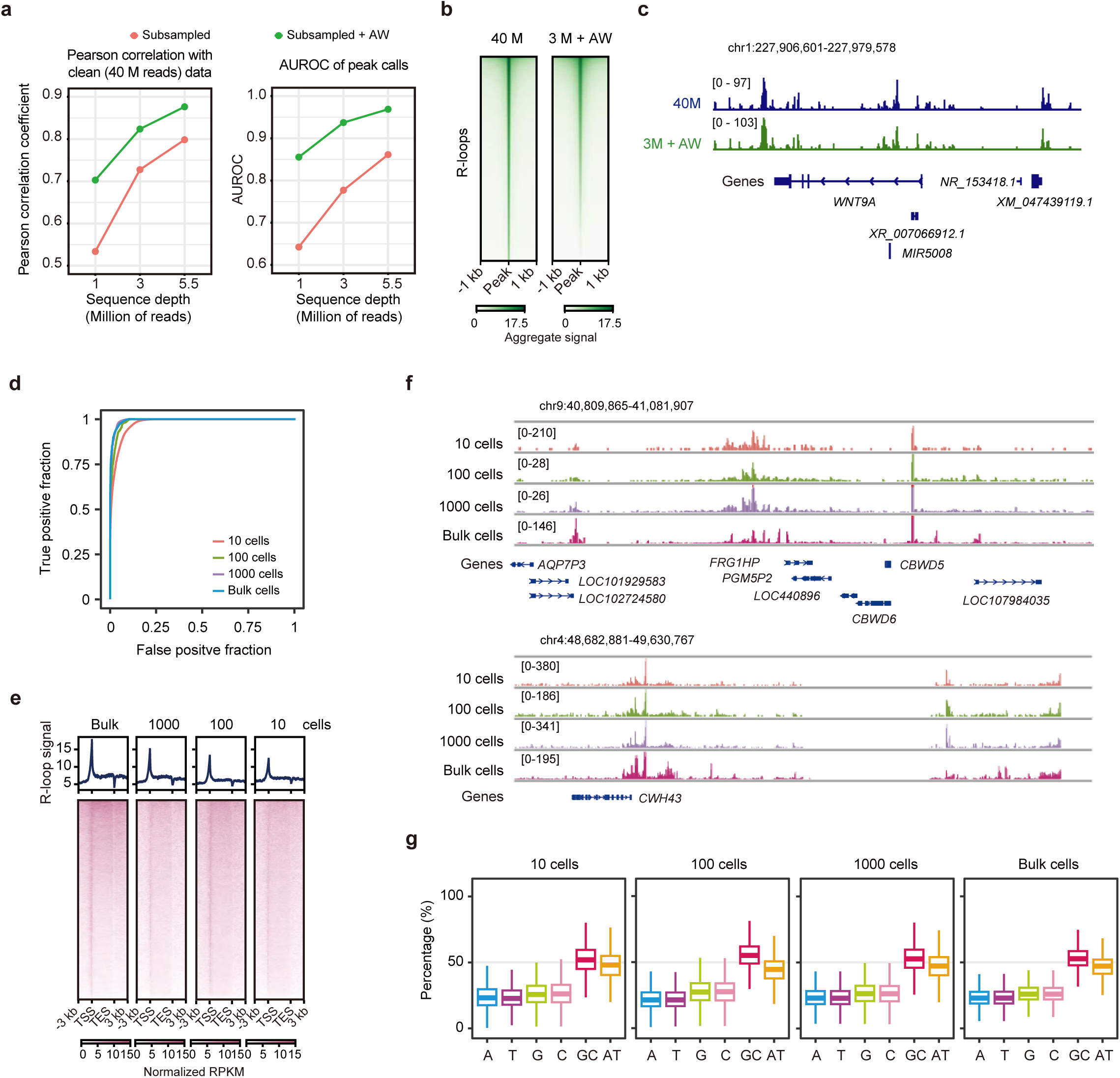
Validation of RIAN-seq across different cell numbers. **a**, Left: Pearson correlation coefficient between 40 million (M) reads dataset and subsampled datasets of RIAN-seq in HEK293T cells with varying sequencing depths before (red) and after (green) AtacWorks (AW) denoising. Right: Area Under the Receiver-Operator Characteristic (AUROC) comparing peak-calling performance of MACS2 (red) and AtacWorks (green) on subsampled data, using MACS2-identified peaks from 40 M reads dataset as a benchmark. **b**, Heatmaps illustrating R-loop signals in 40 M reads dataset and 3 M reads dataset after AW-denoising. **c**, Tracks showing R-loop signals in 40 M reads dataset and 3 M reads dataset with AW-denoising. **d**, Receiver operating characteristic (ROC) curves for RIAN-seq data generated from different numbers of HEK293T cells. **e**, Metaplots and heatmaps depicting R-loop signals (Z-score normalized) obtained from RIAN-seq across gene bodies in HEK293T cells at different input levels (10, 10^2^, 10^3^, and bulk cells). **f**, Tracks showing R-loop enrichment across different numbers of HEK293T cells (10, 10^2^, 10^3^, and bulk cells). **g**, Box plots comparing the base composition of R-loops identified from different numbers of HEK293T cells (10, 10^2^, 10^3^, and bulk cells). Center line, median; box, 25th and 75th percentiles; whiskers, 1.5 × IQR.

**Extended Data Fig. 2.**
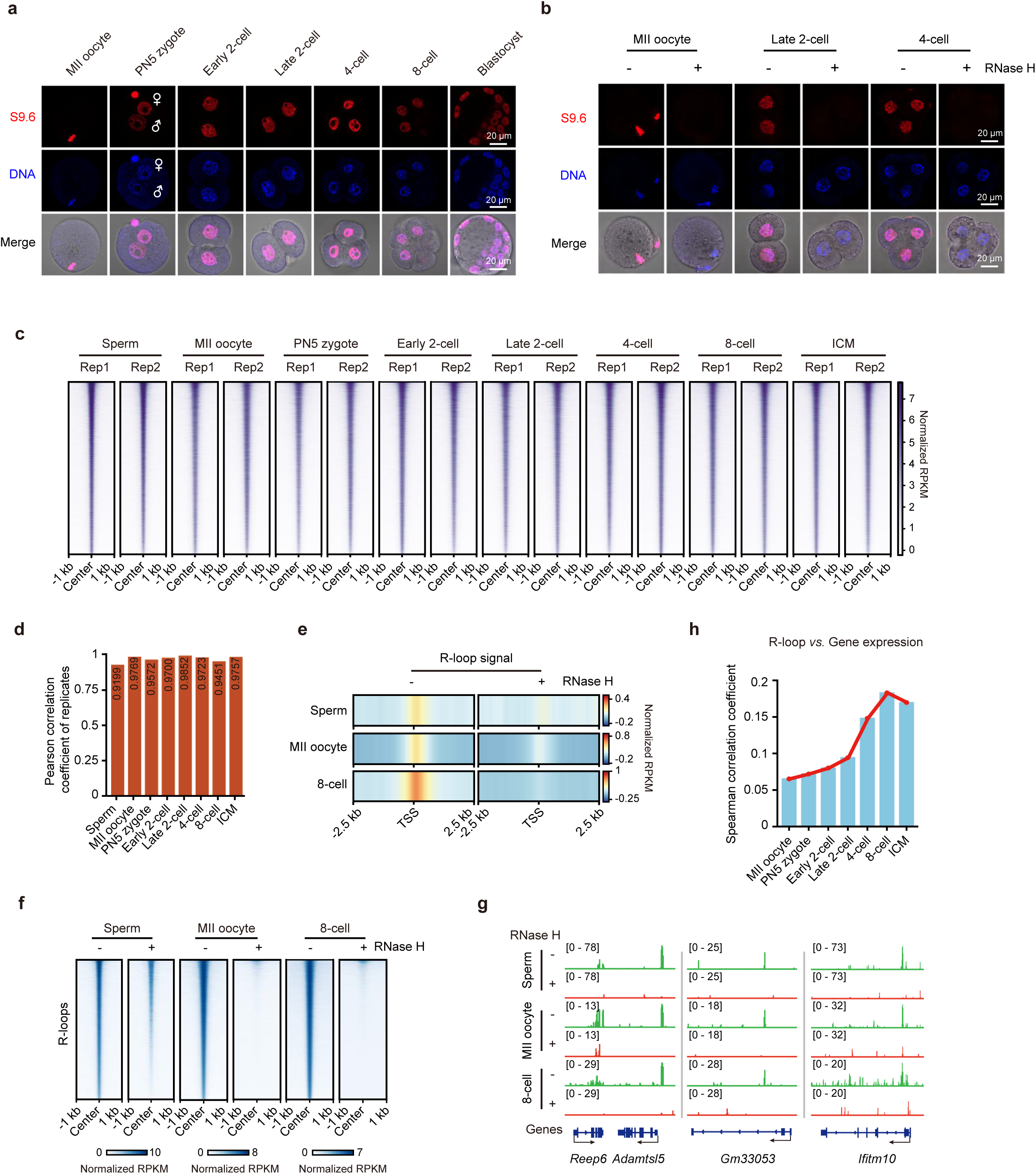
Validation of R-loop datasets in early mouse embryos. **a**, Immunostaining of R-loops (detected using the S9.6 antibody) at various stages of mouse MII oocytes and preimplantation embryos. Sample sizes: MII oocytes (n = 11), PN5 zygotes (n = 12), early 2-cell (n = 10), late 2-cell (n = 12), 4-cell (n = 14), 8-cell (n = 16), and blastocysts (n = 13). A representative image from three independent experiments is shown. Female (♀) and male (♂) pronuclei are labelled. Scale bar: 20 μm. **b**, Immunostaining of R-loops in MII oocytes and preimplantation embryos at late 2-cell and 4-cell stages, with or without RNase H treatment. Sample sizes: MII oocytes (Ctrl, n = 12, RNase H-treated, n = 12), late 2-cell (Ctrl, n = 10, RNase H-treated, n = 10), 4-cell (Ctrl, n = 13, RNase H-treated, n = 10). A representative image from three independent experiments is shown. Scale bar: 20 μm. **c**, Heatmaps displaying R-loop signals (Z-score normalized) across two biological replicates (Rep) in mouse gametes and early embryos at different developmental stages. **d**, Bar graphs illustrating Pearson correlation coefficient for R-loop signals between biological replicates in mouse gametes and early embryos at various developmental stages (two biological replicates). **e**, Heatmaps showing promoter R-loop enrichment (Z-score normalized) in sperm, MII oocytes and 8-cell embryos, with or without RNase H treatment. **f**, Heatmaps depicting R-loop enrichment (Z-score normalized) across the genome in sperm, MII oocytes, and 8-cell embryos with or without RNase H treatment. **g**, Tracks displaying R-loop enrichment at selected loci in sperm, MII oocytes, and 8-cell embryos, with or without RNase H treatment. **h**, Bar graphs presenting Spearman correlation coefficient between gene expression levels and promoter R-loop enrichment in mouse MII oocytes and early embryos at different developmental stages.

**Extended Data Fig. 3.**
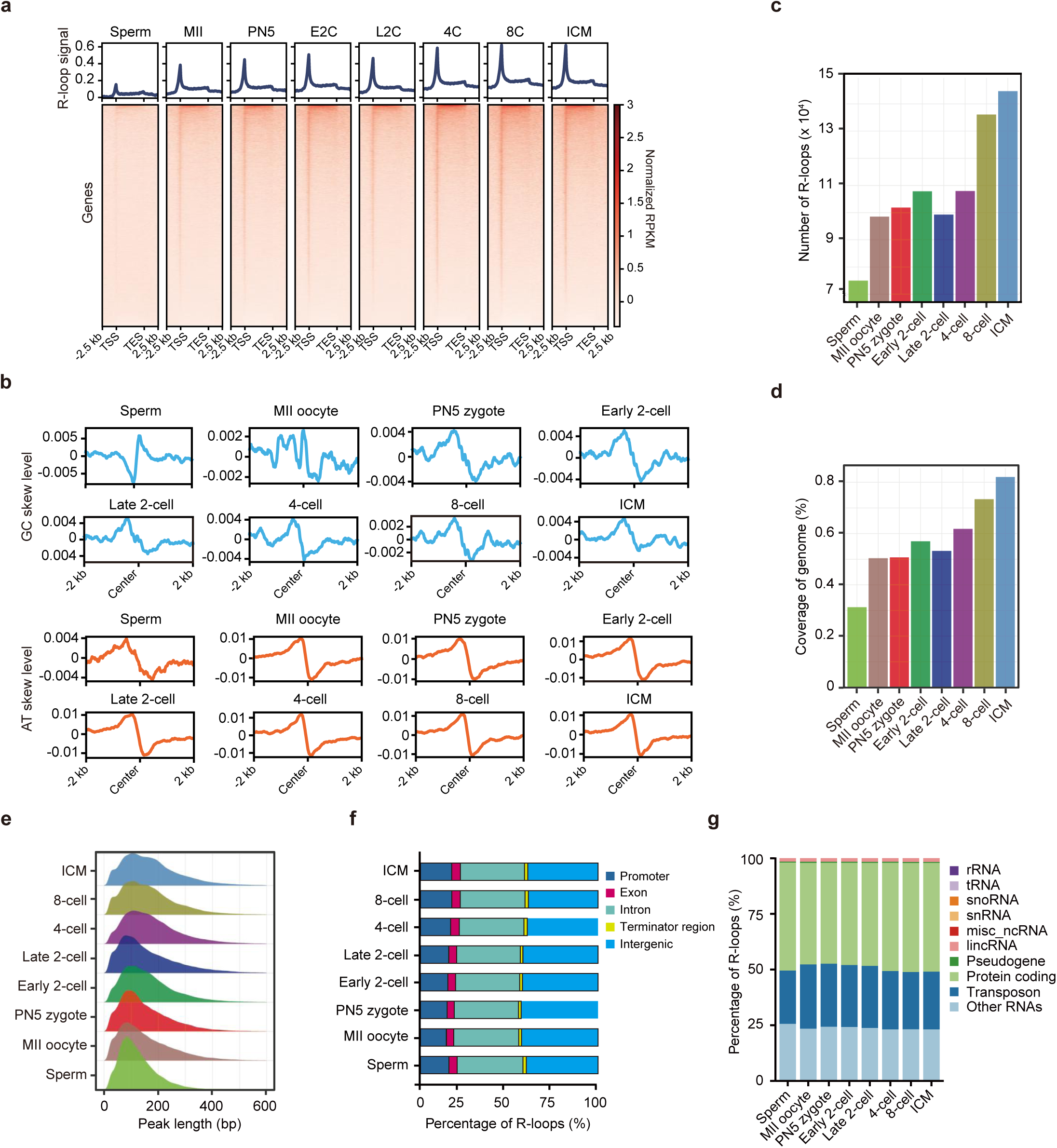
Dynamics of R-loops in mouse gametes and early preimplantation embryos. **a**, Metaplots and heatmaps depicting R-loop signals (Z-score normalized) across genes in mouse gametes and early preimplantation embryos. Developmental stages include sperm, MII oocyte (MII), PN5 zygote (PN5), early 2-cell (E2C), late 2-cell (L2C), 4-cell (4C), 8-cell (8C), inner cell mass from blastocyst (ICM). **b**, Metaplots illustrating GC-and AT-skew levels centered on R-loops of mouse gametes and early preimplantation embryos. **c-d**, Bar graphs representing the total numbers (**c**) and the genomic coverage (**d**) of R-loops detected in mouse gametes and early preimplantation embryos. **e**, Ridgeline plots displaying the length distribution of R-loops in mouse gametes and early preimplantation embryos. **f**, Bar graphs showing the genomic distribution of R-loops across different genomic regions in mouse gametes and early preimplantation embryos. **g**, Bar graphs illustrating the distribution of R-loops at genomic loci associated with different types of RNA transcripts in mouse gametes and early preimplantation embryos.

**Extended Data Fig. 4.**
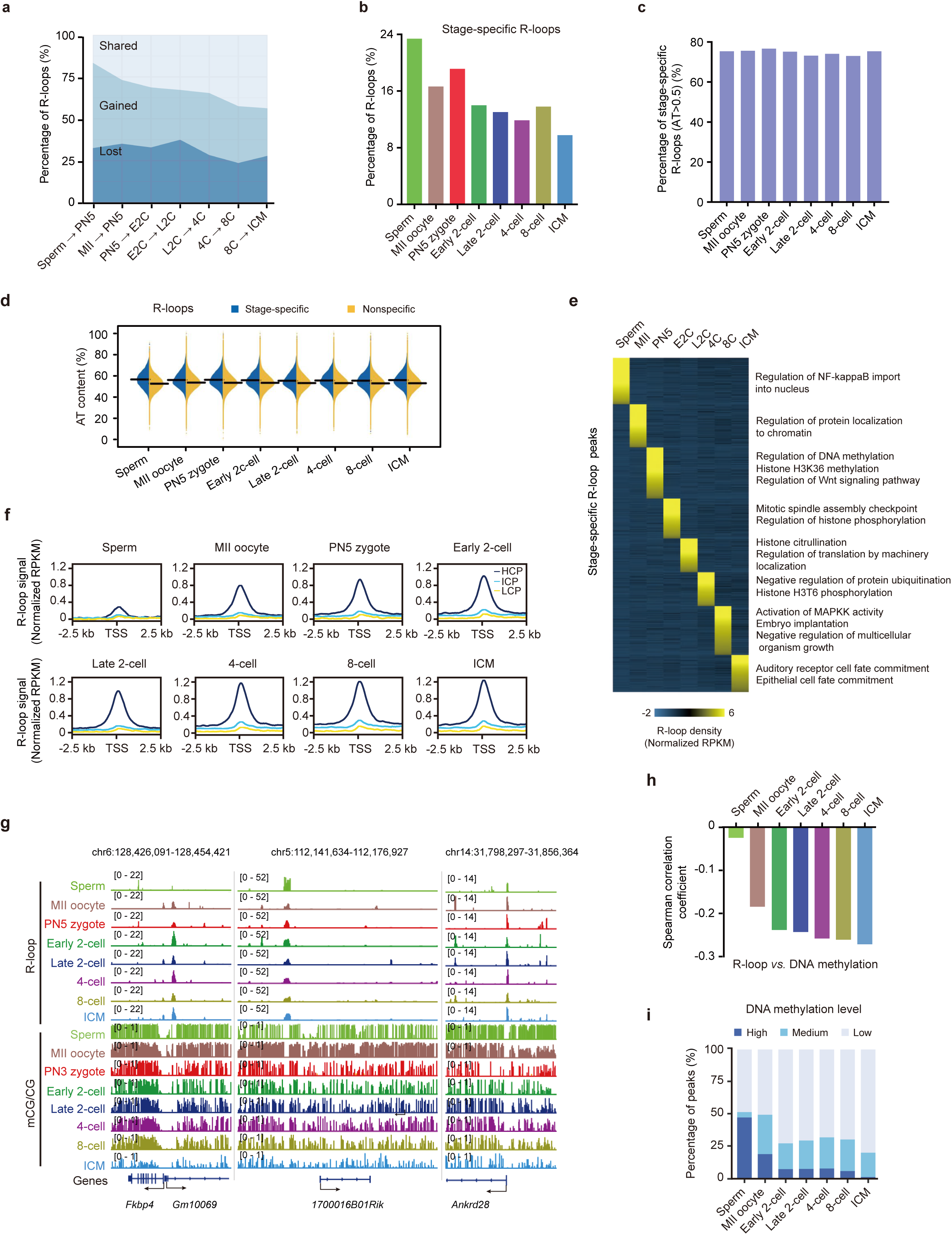
Dynamics of stage-specific R-loops and DNA methylation at R-loops during early preimplantation embryos. **a**, Area charts depicting the percentage of gained, lost, and shared R-loops during transitions between adjacent stages of fertilization and early embryonic development. MII, MII oocyte; PN5, PN5 zygote; E2C, early 2-cell; L2C, late 2-cell; 4C, 4-cell; 8C, 8-cell; ICM, inner cell mass from the blastocyst. **b**, Bar graphs illustrating the proportion of stage-specific R-loops in mouse gametes and early preimplantation embryos. **c**, Bar graphs showing the percentage of AT-rich R-loops within stage-specific R-loops in mouse gametes and early preimplantation embryos. **d**, Bean plots depicting the AT content of stage-specific and nonspecific R-loops in mouse gametes and early preimplantation embryos. The black line represents the median values. **e**, Heatmaps presenting the R-loop enrichment (Z-score normalized) at stage-specific R-loops in mouse gametes and early preimplantation embryos (left). The functional enrichment of genes nearby (by the GREAT analysis) is also shown (right). **f**, Metaplots displaying average R-loop enrichment (Z-score normalized) at high, intermediate, and low CpG promoters (HCPs, ICPs, and LCPs) in mouse gametes and early preimplantation embryos. **g**, Tracks illustrating R-loop enrichment and DNA methylation levels at the selected loci in mouse gametes and early preimplantation embryos. **h**, Bar graphs showing the Spearman correlation coefficient between R-loop enrichment and DNA methylation levels in mouse gametes and early preimplantation embryos. **i**, Bar graphs representing the proportion of R-loops with high (mCG/CG ≥ 0.8), medium (0.2 < mCG/CG < 0.8), and low (mCG/CG ≤ 0.2) DNA methylation levels in mouse gametes and early preimplantation embryos.

**Extended Data Fig. 5.**
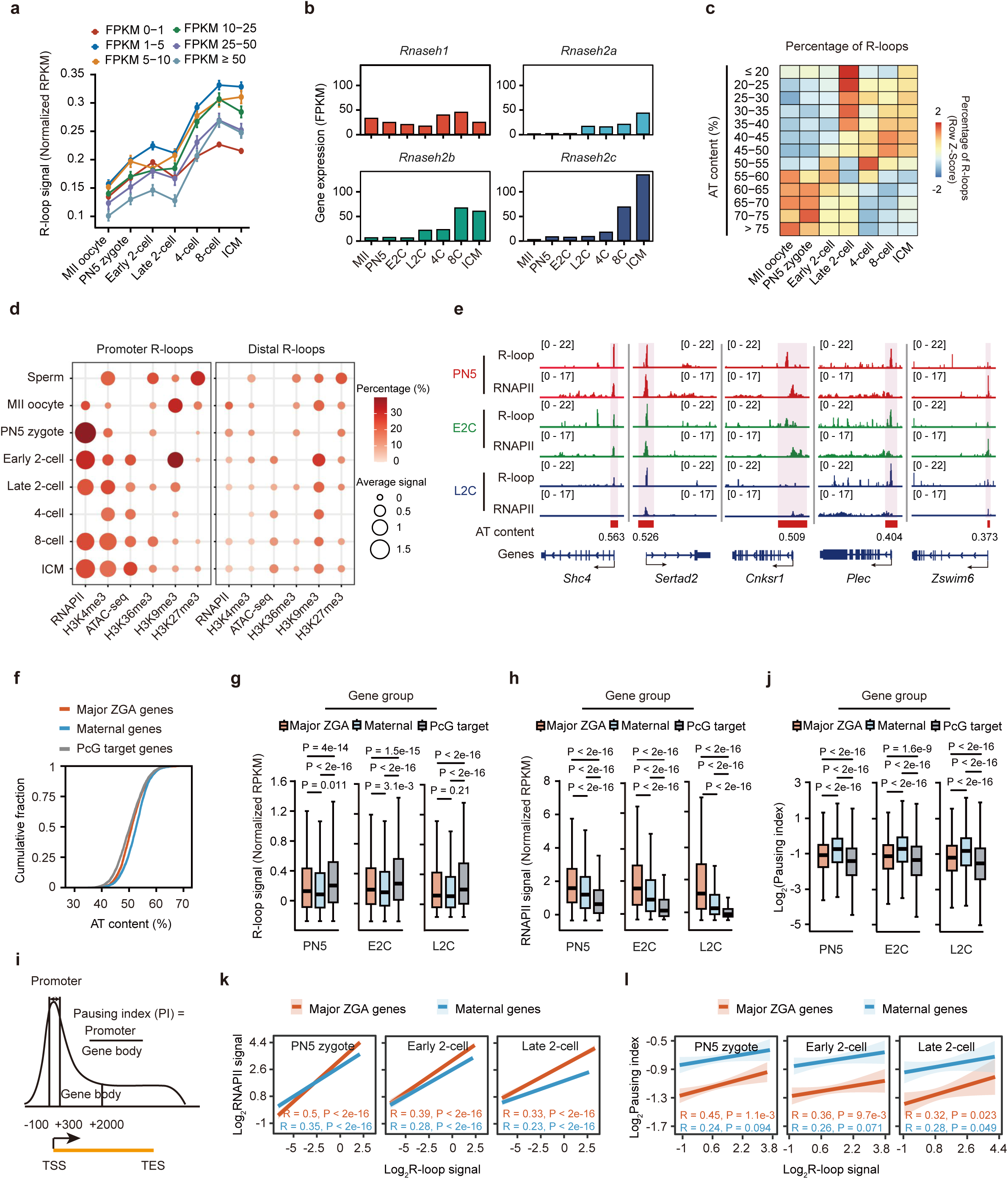
Relationship between R-loops, gene expression, and RNAPII at promoters during early mouse embryonic development. **a**, Line charts depicting the promoter R-loop enrichment (Z-score normalized) with varying expression levels. Error bars, mean ± SE. **b**, Bar graphs illustrating the expression levels of RNase H family members (*Rnaseh1*, *Rnaseh2a*, *Rnaseh2b*, *Rnaseh2c*). **c**, Heatmaps displaying the percentage of R-loops with different AT content. **d**, Enrichment analysis of promoter and distal R-loops in relation to key chromatin features, including RNAPII occupancy, H3K4me3, H3K36me3, H3K27me3, H3K9me3, and chromatin accessibility (ATAC-seq). Bubble size, the average signal intensity (Z-score normalized) of the indicated chromatin features at R-loops. Color gradient, the proportion of R-loops exhibiting significant enrichment (normalized RPKM > 1) of the corresponding chromatin feature. **e**, Tracks showing R-loop and RNAPII enrichment at the promoters with different AT content. Corresponding AT content is indicated. **f**, Cumulative fraction showing the promoter AT content of major ZGA, maternal, and PcG target genes. **g-h**, Box plots comparing promoter R-loop (**g**) and RNAPII (**h**) enrichment (Z-score normalized) of major ZGA, maternal, and PcG target genes. P values, two-sided Wilcoxon rank-sum test. Center line, median; box, 25th and 75th percentiles; whiskers, 1.5 × IQR. **i**, Schematic model showing the definition of the RNAPII pausing index. **j**, Box plots illustrating the RNAPII pausing index of major ZGA, maternal, and PcG target genes during ZGA. P values, two-sided Wilcoxon rank-sum test. **k**, Pearson correlation between promoter R-loop and RNAPII enrichment of major ZGA and maternal genes during ZGA. **l**, Spearman correlation between promoter R-loop enrichment and the RNAPII pausing index of major ZGA and maternal gene during ZGA.

**Extended Data Fig. 6.**
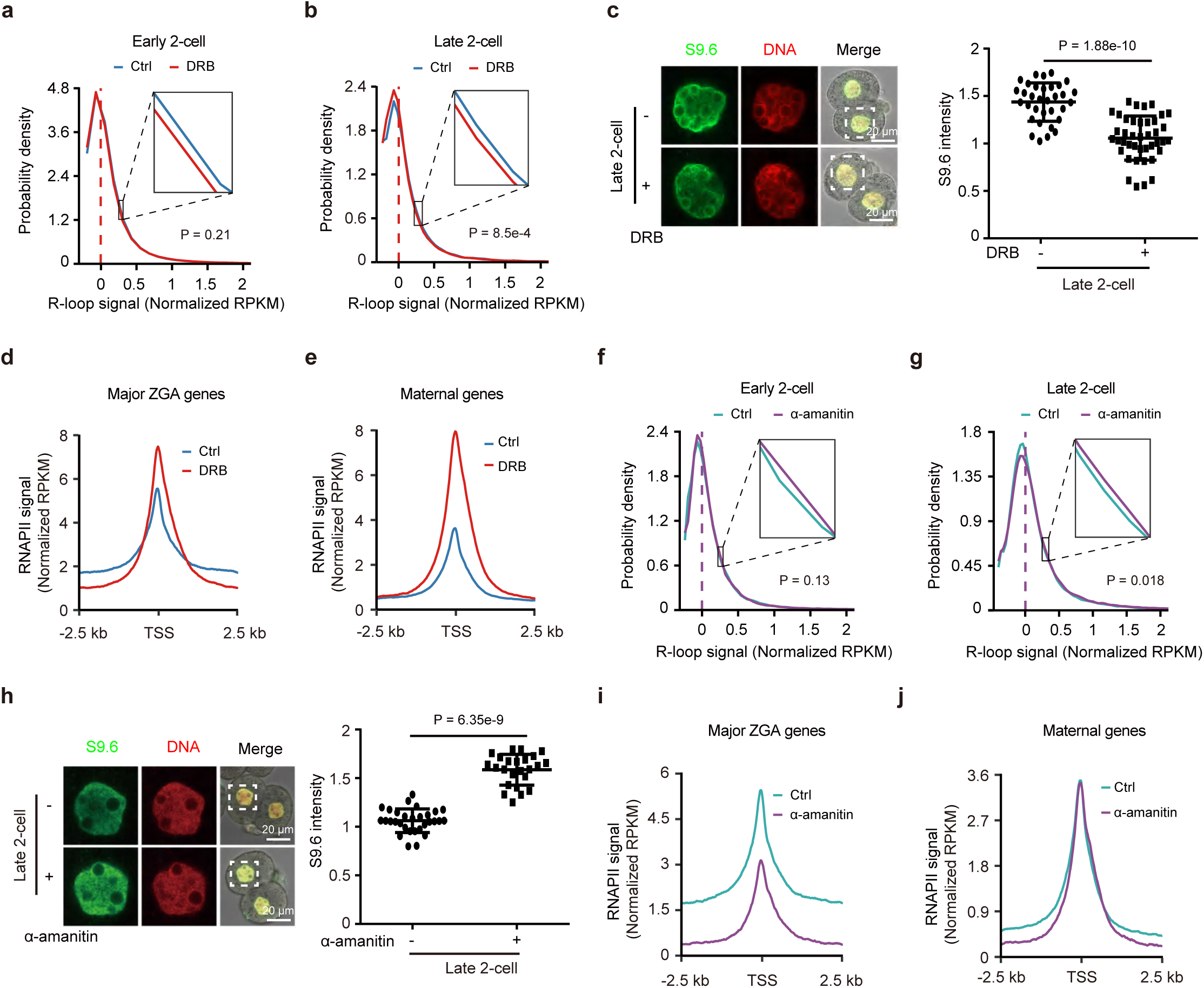
R-loops are independent of RNAPII co-transcription at promoters during ZGA. **a-b**, Probability density plots showing the distribution of global R-loop signals (Z-score normalized) in embryos with or without DRB treatment at early 2-cell (**a**) and late 2-cell (**b**) stages. P values, two-sided Wilcoxon rank-sum test. Dashed line, median values. **c**, Immunostaining of R-loop signals in embryos with or without DRB treatment at late 2-cell stage (Control, n = 17; DRB, n = 22). A representative image from three independent experiments is shown. The dashed squares indicate the nucleus. Scale bar: 20 μm. Quantification of R-loop intensity and P value (two-sided *t*-test) are shown (right). Error bars, mean ± SE. **d-e**, Metaplots showing the average RNAPII enrichment (Z-score normalized) at TSSs of major ZGA (**d**) and maternal genes (**e**) in embryos with or without DRB treatment at late 2-cell stage. **f-g**, Probability density plots illustrating the distribution of R-loop signals (Z-score normalized) in embryos with or without α-amanitin treatment at early 2-cell (**f**) and late 2-cell (**g**) stages. P values, two-sided Wilcoxon rank-sum test. Dashed line, median values. **h,** Immunostaining analysis of R-loop signals in embryos with or without α-amanitin treatment at late 2-cell stage. (Control, n = 15; α-amanitin, n = 13). A representative image from three independent experiments is shown. The dashed squares indicate the nucleus. Scale bar: 20 μm. Quantification of R-loop intensity and P value (two-sided *t*-test) are shown (right). Error bars, mean± SE. **i-j**, Metaplots showing the average RNAPII enrichment (Z-score normalized) at TSSs of major ZGA (**i**) and maternal genes (**j**) in embryos with or without α-amanitin treatment at late 2-cell stage.

**Extended Data Fig. 7.**
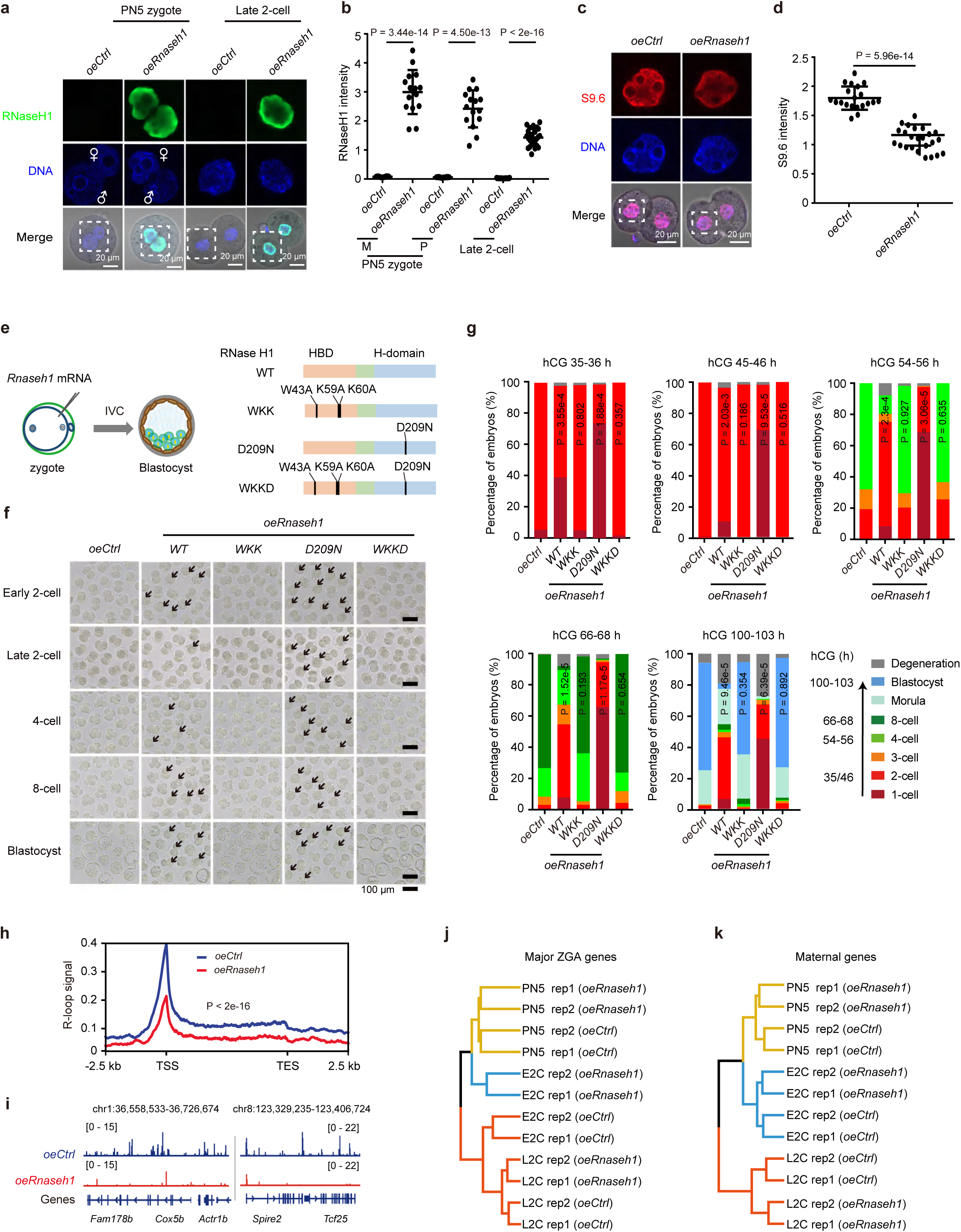
The loss of R-loops impairs embryonic development during ZGA and affects gene expression. **a-b**, Immunostaining (**a**) and quantifications (**b**) of RNase H1 in control (*oeCtrl*) and *Rnaseh1-*overexpressed (*oeRNaseh1*) PN5 zygotes (*oeCtrl*, n = 14; *oeRnaseh1*, n = 15) and late 2-cell embryos (*oeCtrl*, n = 19; *oeRnaseh1*, n = 13). One representative image from three independent experiments is shown. Female (♀) and male (♂) pronuclei are labelled. The dashed squares indicate the nucleus. Scale bar: 20 μm. P values, two-sided *t*-test. Error bars, mean ± SE. M, female pronucleus; P, male pronucleus. **c-d**, Immunostaining (**c**) and quantifications (**d**) of R-loop intensity in control (*oeCtrl*) and *Rnaseh1-*overexpressed (*oeRnaseh1*) late 2-cell embryos (*oeCtrl*, n = 10; *oeRnaseh1*, n = 12). One representative image from three independent experiments is shown. The dashed squares indicate the nucleus. Scale bar, 20 μm. P value, two-sided *t*-test. Error bars, mean ± SE. **e**, Schematic representation of microinjection of *Rnaseh1* (*WT* and *mutant*) into early embryos and *in vitro* culture with KSOM medium. **f-g**, Embryo morphology (**f**) and developmental rates (**g**) of control (*oeCtrl*) and *Rnaseh1* (WT and mutants) overexpressed embryos (*oeCtrl*, n = 40, 40, 53, 42, 49, 35; *oeRnaseh1* (*WT*), n = 40, 55, 41; *oeRnaseh1* (*WKK*), n = 41, 33, 52; *oeRnaseh1* (*D209N*), n = 59, 53, 45); *oeRnaseh1* (*WKKD*), n = 40, 42, 52). Arrows, abnormal embryos. Scale bar, 100 μm. P values, two-sided *t-*test. **h**, Metaplots comparing R-loop enrichment (Z-score normalized) between control (*oeCtrl*) and *Rnaseh1*-overexpressed (*oeRnaseh1*) late 2-cell embryos. P value, two-sided Wilcoxon rank-sum test. **i**, Tracks showing R-loop enrichment at selected loci in control (*oeCtrl*) and *Rnaseh1-*overexpressed (*oeRnaseh1*) late 2-cell embryos. **j-k**, Hierarchical clustering analysis showing the expression of major ZGA (**j**) and maternal genes (**k**) in control (*oeCtrl*) and *Rnaseh1-*overexpressed (*oeRnaseh1*) embryos (two biological replicates, Rep). PN5, PN5 zygote; E2C, early 2-cell; L2C, late 2-cell.

**Extended Data Fig. 8.**
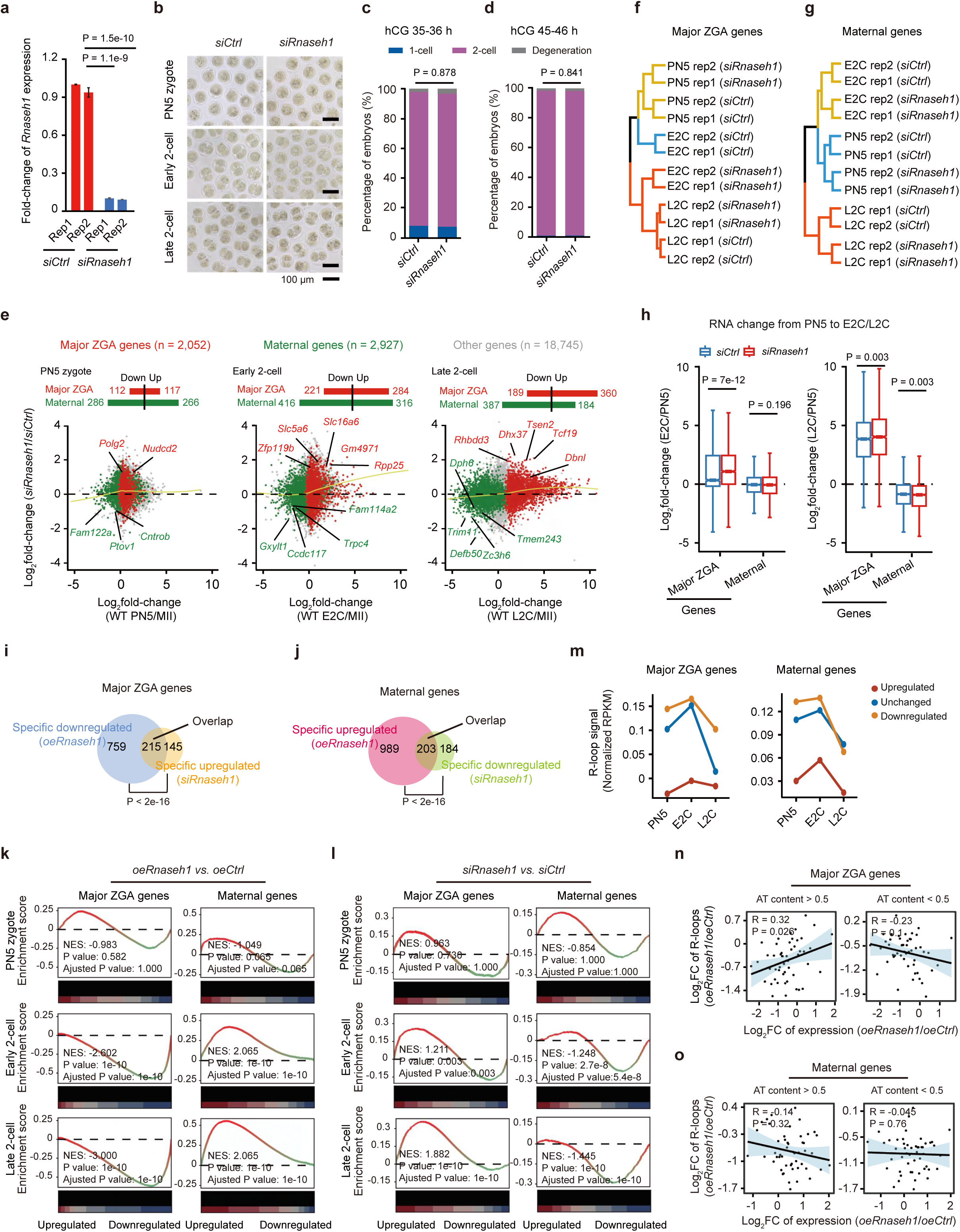
The effect of *Rnaseh1* knockdown on the expression of major ZGA and maternal genes. **a**, Bar graphs showing the *Rnaseh1* expression in control (*siCtrl*) and *Rnaseh1-*knockdown (*siRnaseh1*) late 2-cell embryos (two biological replicates, Rep). P values, two-sided *t*-test. Error bars, mean ± SD **b-d**, Embryo morphology (**b**) and developmental rate (**c**, **d**) of control (*siCtrl*) and *Rnaseh1-*knockdown (*siRnaseh1*) embryos. Scale bars: 100 μm. Early 2-cell (hCG 35-36 h) (*siCtrl*, n = 42, 45, 46; *siRnaseh1*, n = 43, 36, 42) (**c**); late 2-cell (hCG 45-46 h) (*siCtrl*, n = 42, 45, 46; *siRnaseh1*, n = 43, 36, 42) (**d**). P values, two-sided *t*-test. **e**, Scatter plots showing gene expression fold-changes upon *Rnaseh1* knockdown (two biological replicates). Yellow lines, local regression fitting. **f-g**, Hierarchical clustering analysis showing expression of major ZGA (**f**) and maternal genes (**g**) in control (*siCtrl*) or *Rnaseh1*-knockdown (*siRnaseh1*) embryos (two biological replicates, Rep). PN5, PN5 zygote; E2C, early 2-cell; L2C, late 2-cell. **h**, Box plots showing expression changes of major ZGA and maternal genes from PN5 zygote to early 2-cell or late 2-cell stage in control (*siCtrl*) and *Rnaseh1*-knockdown (*siRnaseh1*) embryos (two biological replicates). P values, two-sided Wilcoxon rank-sum test. Center line, median; box, 25th and 75th percentiles; whiskers, 1.5 × IQR. **i**, Venn diagram showing the overlap between downregulated major ZGA genes upon *Rnaseh1* overexpression (*oeRnaseh1*) and upregulated major ZGA genes upon *Rnaseh1* knockdown (*siRnaseh1*) at late 2-cell stage. P value, Fisher’s exact test. **j**, Venn diagram showing the overlap between upregulated maternal genes upon *Rnaseh1* overexpression (*oeRnaseh1*) and downregulated maternal genes upon *Rnaseh1* knockdown (*siRnaseh1*) at late 2-cell stage. P value, Fisher’s exact test. **k-l**, Gene set enrichment analysis (GSEA) of major ZGA and maternal genes in embryos upon *Rnaseh1* overexpression (*oeRnaseh1*) (**k**) or *Rnaseh1* knockdown (*siRnaseh1*) (**l**). NES, normalized enrichment score. **m**, Line charts representing promoter R-loop enrichment (Z-score normalized) of upregulated, unchanged, and downregulated major ZGA (left) and maternal genes (right) after *Rnaseh1* overexpression in normal embryos during ZGA. Data are presented as median values. **n-o**, Scatter plots showing the Spearman correlation between fold-change (FC) in gene expression and fold-change in promoter R-loop enrichment of major ZGA (**n**) and maternal genes (**o**) after *Rnaseh1* overexpression, categorized by AT content of promoter R-loops (major ZGA genes, AT > 0.5, n = 789; AT < 0, n = 1,261; maternal genes, AT > 0.5, n = 1,104; AT < 0, n = 1,526). Each group of genes was categorized into 50 bins based on their ranked fold-change in expression at late 2-cell stage upon *Rnaseh1* overexpression (*oeRnaseh1*).

**Extended Data Fig. 9.**
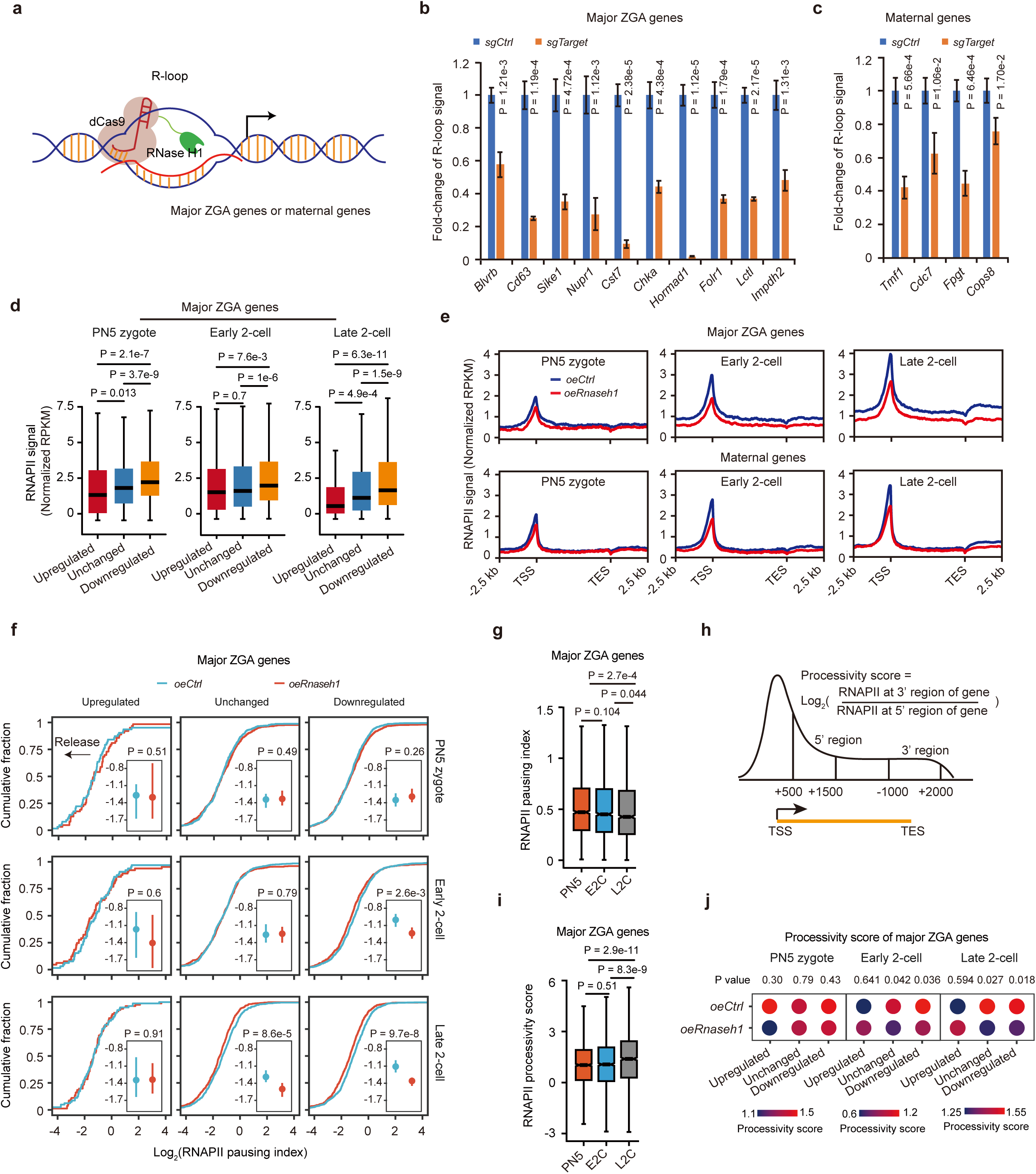
The effect of R-loop loss on RNAPII. **a**, Diagram of site-specific regulation of R-loops. *dCas9*-*Rnaseh1* mRNA and gRNA targeting specific R-loops were microinjected at 22-23 h after hCG treatment, and the embryos were then cultured *in vitro*. **b-c**, Bar graphs showing promoter R-loop enrichment of selected major ZGA (**b**) and maternal genes (**c**) upon promoter AT-rich R-loop loss. P values, two-sided *t*-test. Error bars, mean ± SD. **d**, Box plots showing promoter RNAPII enrichment (Z-score normalized) in normal embryos of upregulated, unchanged, and downregulated major ZGA genes after *Rnaseh1* overexpression. P values, two-sided Wilcoxon rank-sum test. Center line, median; box, 25th and 75th percentiles; whiskers, 1.5 × IQR. **e**, Metaplots showing the average RNAPII enrichment (Z-score normalized) across major ZGA (top) and maternal genes (bottom) in control (*oeCtrl*) and *Rnaseh1*-overexpressed (*oeRnaseh1*) embryos. **f**, Cumulative distribution and error bar plots (insert) showing the RNAPII pausing index for upregulated, unchanged, and downregulated major ZGA genes in control (*oeCtrl*) and *Rnaseh1*-overexpressed (*oeRnaseh1*) embryos. P values, two-sided Wilcoxon rank-sum test. Data are presented as median ± 95% CIs in the error bar plots. **g**, Box plots showing the RNAPII pausing index on major ZGA genes in normal embryos. P values, two-sided Wilcoxon rank-sum test. Center line, median; box, 25th and 75th percentiles; whiskers, 1.5 × IQR. **h**, Schematic model showing the definition of the RNAPII processivity score. **i**, Box plots showing the RNAPII processivity score on major ZGA genes in normal embryos. P values, two-sided Wilcoxon rank-sum test. Center line, median; box, 25th and 75th percentiles; whiskers, 1.5 × IQR. **j**, Ballon plots showing the average RNAPII processivity scores on upregulated, unchanged, and downregulated major ZGA genes in control (*oeCtrl*) and *Rnaseh1*-overexpressed (*oeRnaseh1*) embryos. P values, two-sided *t*-test. Data are presented as mean values.

**Extended Data Fig. 10.**
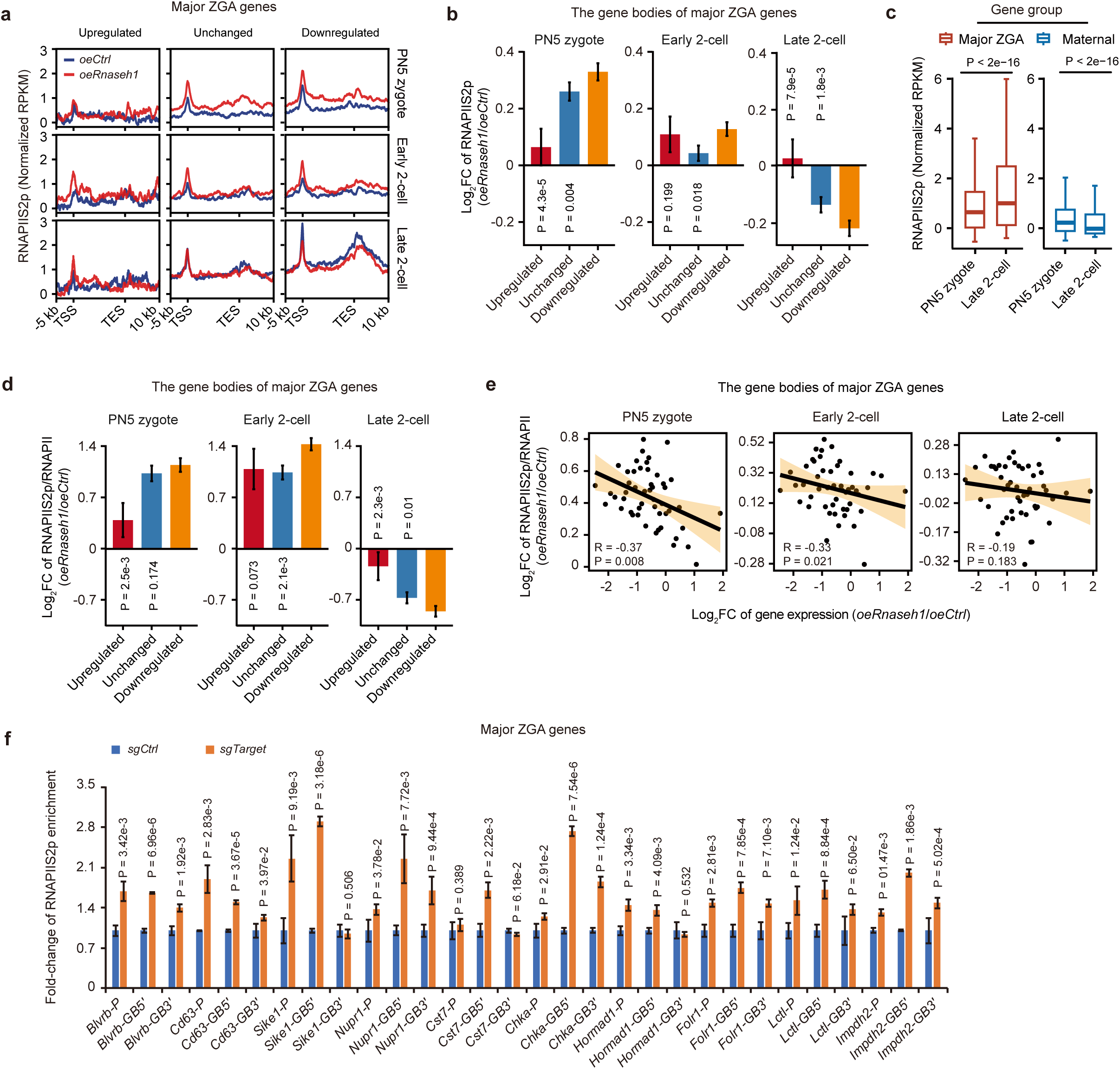
R-loop loss induces RNAPII pause release on and major ZGA and maternal genes during ZGA. **a**, Metaplots showing the average RNAPIIS2p enrichment (Z-score normalized) across upregulated, unchanged, and downregulated major ZGA genes after *Rnaseh1* overexpression. **b**, Bar graphs showing the fold-change (FC) of RNAPIIS2p at the gene bodies (TSS + 0.3 kb to TES) of major ZGA genes upon *Rnaseh1* overexpression. Each group is compared to downregulated major ZGA genes. P values, two-sided Wilcoxon rank-sum test. Error bars, mean ± SE. **c**, Box plots showing RNAPIIS2p enrichment (Z-score normalized) at the gene bodies (TSS + 0.3 kb to TES) of major ZGA and maternal genes, in normal PN5 zygote and late 2-cell embryos. P values, two-sided Wilcoxon rank-sum test. Center line, median; box, 25th and 75th percentiles; whiskers, 1.5 × IQR. **d**, Bar graphs showing the fold-change (FC) of RNAPIIS2p/RNAPII ratio at the gene bodies (TSS + 0.3 kb to TES) of major ZGA genes upon *Rnaseh1* overexpression. Each group is compared to downregulated major ZGA genes. P values, two-sided Wilcoxon rank-sum test. Error bars, mean ± SE. **e**, Scatter plots demonstrating the Spearman correlation between the fold-change (FC) of expression and gene body RNAPIIS2p/RNAPII ratio (TSS + 0.3 kb to TES) in major ZGA genes at late 2-cell embryos upon *Rnaseh1* overexpression during ZGA. Genes were categorized into 50 bins based on their ranked fold-change of expression at late 2-cell stage upon *Rnaseh1* overexpression. **f**, Bar graphs illustrating the relative enrichment of RNAPIIS2p at the promoters and gene bodies of selected major ZGA genes in early 2-cell embryos following the loss of promoter AT-rich R-loops. P values, two-sided *t*-test. Error bars, mean ± SD.

**Extended Data Fig. 11.**
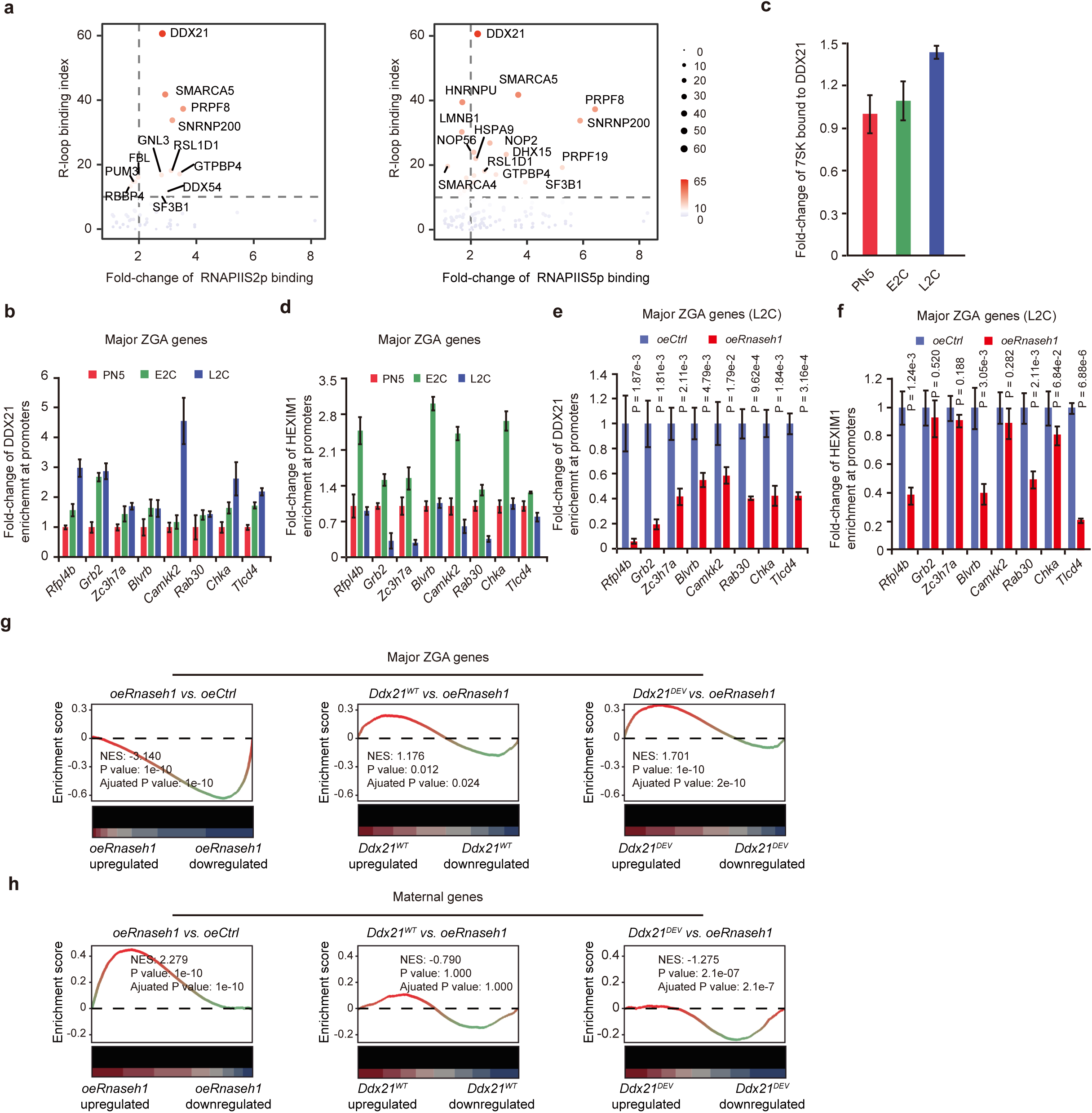
R-loop loss Promote CDK9 release from 7SK/HEXIM1snRNP complex on major ZGA and maternal genes during ZGA. **a**, Volcano plots showing significantly enriched factors from RNAPIIS2p (left) or RNAPIIS5p (right) interactomes, overlapped with the R-loop interactome in mES cells. **b**, Bar graphs showing the relative enrichment of DDX21 at the promoters of selected major ZGA genes in normal embryos. P values, two-sided *t*-test. Error bars, mean ± SD. PN5, PN5 zygote; E2C, early 2-cell; L2C, late 2-cell. **c**, Bar graphs showing the relative enrichment of 7SK bound to DDX21 in normal embryos. P values, two-sided *t*-test. Error bars, mean ± SD. **d**, Bar graphs showing the relative enrichment of HEXIM1 at the promoters of selected major ZGA genes in normal embryos. P values, two-sided *t*-test. Error bar mean ± SD. **e**, Bar graphs showing the relative enrichment of DDX21 enrichment at the promoters of selected major ZGA genes in control (*oeCtrl*) and *Rnaseh1*-overexpressed (*oeRnaseh1*) embryos at late 2-cell (L2C) stage. P values, two-sided *t*-test. Error bars, mean ± SD. **f**, Bar graphs depicting the relative enrichment in HEXIM1 enrichment at the promoters of selected major ZGA genes in control (*oeCtrl*) and *Rnaseh1*-overexpressed (*oeRnaseh1*) embryos at late 2-cell (L2C) stage. P values, two-sided *t*-test. Error bars, mean ± SD. **g, h**, GSEA of major ZGA (**g**) and maternal genes (**h**) in embryos with *Rnaseh1* overexpression (*oeRnaseh1*), *Rnaseh1* overexpression combined with endogenous *Ddx21* knockdown, rescued by either wild-type *Ddx21* (*Ddx21^WT^*) or a helicase activity defective mutant (*Ddx21^DEV^*). NES, normalized enrichment score.

**Extended Data Fig. 12.**
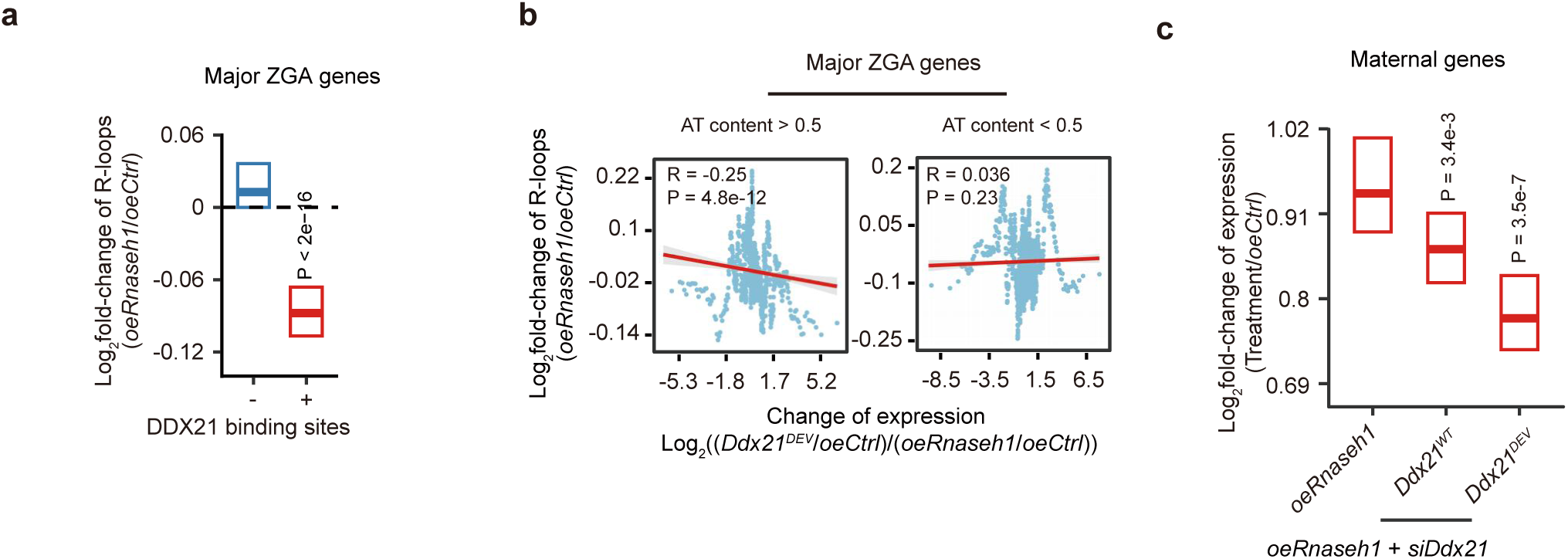
Helicase-deficient DDX21 rescues the expression of major ZGA and maternal genes following R-loop loss. **a**, Crossbar plots showing the promoter R-loop enrichment fold-change for major ZGA genes, with or without the DDX21 binding sites, after *Rnaseh1* overexpression. P values, two-sided Wilcoxon rank-sum test. Data are presented as median ± 95% CIs. **b**, Scatter plots displaying the Spearman correlation between the fold-change of expression and promoter R-loop enrichment fold-change in major ZGA genes after *Rnaseh1* overexpression (*oeRNaseh1*), *Rnaseh1* overexpression combined with endogenous *Ddx21* knockdown, rescued by either wild-type *Ddx21* (*Ddx21^WT^*) or a helicase-defective mutant (*Ddx21^DEV^*). Data were analyzed using a moving window approach (window size = 25 genes, step = 1 gene). Genes were classified based on the AT content of promoter R-loops (AT > 0.5, n = 789; AT < 0.5, n = 1,261). **c**, Crossbar plots showing the fold-change of expression in maternal genes after *Rnaseh1* overexpression (*oeRnaseh1*), *Rnaseh1* overexpression combined with endogenous *Ddx21* knockdown, rescued by wild-type *Ddx21* (*Ddx21^WT^*) or a helicase-defective mutant (*Ddx21^DEV^*), relative to control. Genes were categorized by with or without DDX21 binding sites. P values, two-sided Wilcoxon rank-sum test. Data are presented as median ± 95% CIs.

